# Annual phytoplankton dynamics in coastal waters from Fildes Bay, Western Antarctic Peninsula

**DOI:** 10.1101/2020.10.27.356600

**Authors:** Nicole Trefault, Rodrigo De la Iglesia, Mario Moreno-Pino, Adriana Lopes dos Santos, Catherine Gérikas Ribeiro, Antonia Cristi, Dominique Marie, Daniel Vaulot

## Abstract

Year-round reports of phytoplankton dynamics in the West Antarctic Peninsula are rare and mainly limited to microscopy and/or pigment-based studies. We analyzed the phytoplankton community from coastal waters of Fildes Bay in the West Antarctic Peninsula between January 2014 and 2015 using metabarcoding of the nuclear and plastidial 18/16S rRNA gene from both size-fractionated and flow cytometry sorted samples. Each metabarcoding approach yielded a different image of the phytoplankton community with for example Prymnesiophyceae more prevalent in plastidial metabarcodes and Mamiellophyceae in nuclear ones. Overall 14 classes of photosynthetic eukaryotes were present in our samples with the following dominating: Bacillariophyta (diatoms), Pelagophyceae and Dictyochophyceae for division Ochrophyta, Mamiellophyceae and Pyramimonadophyceae for division Chlorophyta, Prymnesiophyceae and Cryptophyceae. Diatoms were dominant in the larger size fractions and during summer, while Prymnesiophyceae and Cryptophyceae were dominant in colder seasons. Pelagophyceae were particularly abundant towards the end of autumn (May). In addition of *Micromonas polaris* and *Micromonas* sp. clade B3, both previously reported in Arctic waters, we detected a new *Micromonas* 18S rRNA sequence signature, close to but clearly distinct from *M. polaris*, which potentially represent a new clade specific of the Antarctic. These results highlight the need for complementary strategies as well as the importance of year-round monitoring for a comprehensive description of phytoplankton communities in Antarctic coastal waters.

## Introduction

Phytoplankton represents the main energy input to the marine ecosystem in Antarctica, providing fixed carbon to marine and terrestrial systems, being the primary food source, and therefore the base of the entire Antarctic food web (Browning et al. 2014; Smetacek and Nicol 2005). Summer phytoplankton blooms in nutrient rich coastal waters are critical to fuel the Antarctic marine ecosystem and to maintain energy fluxes during the long winter. Each year, the temperature increase and the melting of ice during the Austral spring induces a succession of phytoplankton communities which understanding is crucial, since it has profound implications at planetary scales, from the architecture and efficiency of the trophic webs, to the carbon sedimentation to deep waters and the global biogeochemical cycles (Garibotti et al. 2005). Monitoring natural phytoplankton populations is challenging, especially in high latitude environments such Antarctica given logistical field difficulties. Long time series such as the Rothera Time Series (RaTS) and the Palmer Long-Term Ecological Research (PAL-LTER) program help understanding of the year-round Antarctic phytoplankton dynamics.

The Western Antarctic Peninsula (WAP) is one of the fastest warming areas on Earth (Clem et al. 2020) and is characterized by strong spatial and temporal variability (Martinson et al. 2008). Previous studies have shown regional differences between the northern and southern areas of the WAP, mainly related to mixed layer depth and phytoplankton productivity (Schofield et al. 2018), as well as inter-decadal variability of phytoplankton biomass along the coast of the WAP, with essential role of local-scale forcing on phytoplankton dynamics (Kim et al. 2018). Differences between WAP eastern and western coastal areas have also been described, the former mostly dominated by benthic diatoms and the latter by pelagic ones (Lange et al. 2018). A two year sampling study in Admiralty Bay (King George Island, WAP) reported that spring-summer biomass maxima were dominated by pico-phytoplankton and nanosized flagellates, followed in abundance by diatoms and dinoflagellates (Kopczynska 2008). In Ryder Bay (Adelaide Island), high temperatures were reported to be correlated with increased nano-sized cryptophytes abundance, whereas the haptophyte *Phaeocystis antarctica* increased in relation to high irradiance and low salinity (Biggs et al. 2019). *P. antarctica*, which is replaced by *Phaeocystis pouchetii* in the Arctic ocean (Assmy et al. 2017), is widely present in the WAP (Biggs et al. 2019; Egas et al. 2017) as well as in other Antarctic regions (Arrigo et al. 1999; Delmont et al. 2014). In Fildes Bay (King George Island), phytoplankton showed a rapid increase in biomass and cell abundance as a consequence of short vertical mixing events in the water column, with a strong dominance of nano-phytoplankton, represented by *Thalassiosira* and *Phaeocystis* (Egas et al. 2017). Large diatoms, *Phaeocystis*, and mixotrophic/phagotrophic dinoflagellates, explain most spatial variability in the carbon export potential of the WAP (Lin et al. 2017). More recently, metagenomic and metatranscriptomic analyses of pico- and nano-size fractions of the plankton community from Chile Bay (Greenwich Island, WAP) indicated that while diatoms completely dominated the RNA and DNA-based analyses, alveolates, cryptophytes and haptophytes appear in the RNA-based analyses (possibly corresponding to the active fraction), suggesting that other phytoplankton groups besides diatoms are also actively growing (Alcamán-Arias et al. 2018). From the spatial point of view, variation of phytoplankton across environmental gradients in Fildes Bay, studied using flow cytometry and metabarcoding of the plastidial 16S rRNA gene, indicated, that although the community composition was mostly similar at sub-mesoscale, the abundance of specific phytoplankton groups was responsive to salinity and nutrient inputs (Moreno-Pino et al. 2016).

Environmental sequencing of taxonomic marker genes first by the Sanger technique and then high throughput techniques (metabarcoding) has improved our ability to detect and identify groups that are difficult to cultivate or identify by other methodologies (e.g. microscopy). Two marker genes have been used for phytoplankton diversity studies: nuclear 18S rRNA and plastidial 16S rRNA (Fuller et al. 2006; Moon-van der Staay et al. 2001) yielding quite different images of the community structure (Shi et al. 2011). The use of different cell collection and filtering approaches have also shown differences in the resulting phytoplankton community composition: besides size-fractionation by filtration, a classical approach based on cell size proposed by Sieburth *et al.* (1978), flow cytometry sorting enables to better assess the diversity of small photosynthetic eukaryotes for the pico- and nano-sized fractions (Balzano et al. 2012b; Marie et al. 2010).

In the present study, we sampled the phytoplankton community in coastal waters from Fildes Bay (also called Maxwell Bay, South Shetland Islands, WAP) between January 2014 and 2015, with the aim to assess changes in phytoplankton abundance, diversity and community composition occurring along the Austral year. We used three complementary approaches: size-fractionated samples with nuclear 18S rRNA and plastidial 16S rRNA metabarcoding, and flow cytometry sorted samples with 18S rRNA.

## Results

### Annual phytoplankton variation

We sampled phytoplankton in coastal waters of Fildes Bay, King George Island, at the eastern tip of the WAP (Figure 1A), between January 2014 and 2015 at all seasons except winter (Table 1). Phytoplankton abundance measured by flow cytometry was higher during the summer, compared to the rest of the year (Figure 1B). In autumn, we detected low and uniform levels of the three phytoplankton populations, pico-eukaryotes (PPE), photosynthetic nano-eukaryotes (PNE) and cryptophytes (CRY), with values between 47 and 342 cells mL^−1^ for CRY and PPE, respectively (Supplementary Data S1). CRY showed similar values between summer 2014 and 2015, while PPE and PNE showed an inverted pattern of abundance. PNE were, on average, three times higher than PPE in summer 2015, while it was the reverse in 2014.

**Figure 1.**
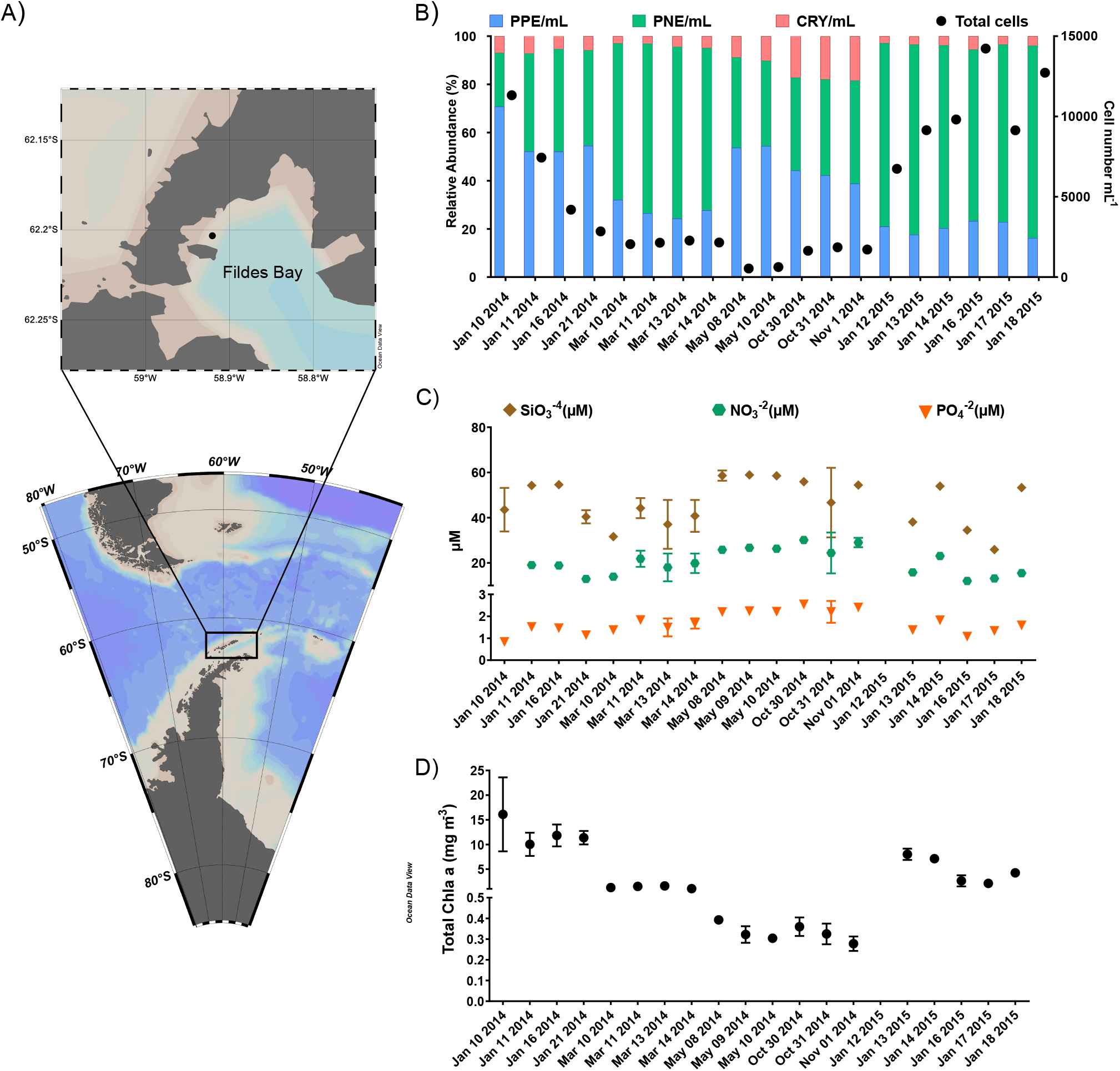
Location of the sampling station in Fildes Bay, King George Island, Western Antarctic Peninsula (WAP) and biotic and abiotic characteristics between January 2014 and January 2015. (A) Map of the Antarctica Peninsula and location of the station in Fildes Bay sampled in this study. (B) Phytoplankton abundance measured by flow cytometry. Detected populations correspond to PPE = photosynthetic pico-eukaryotes, PNE = photosynthetic nano-eukaryotes, and CRY = cryptophytes. (C) Nutrients (silicate, SiO_3_^−2^; nitrate, NO_3_^−^ and phosphate, PO_4_^−3^). (D) Chlorophyll *a* levels during the sampling period. Values correspond to < 100 *μ*m biomass. For B, C, and D, values represent mean ± standard deviation.

**Table 1.**
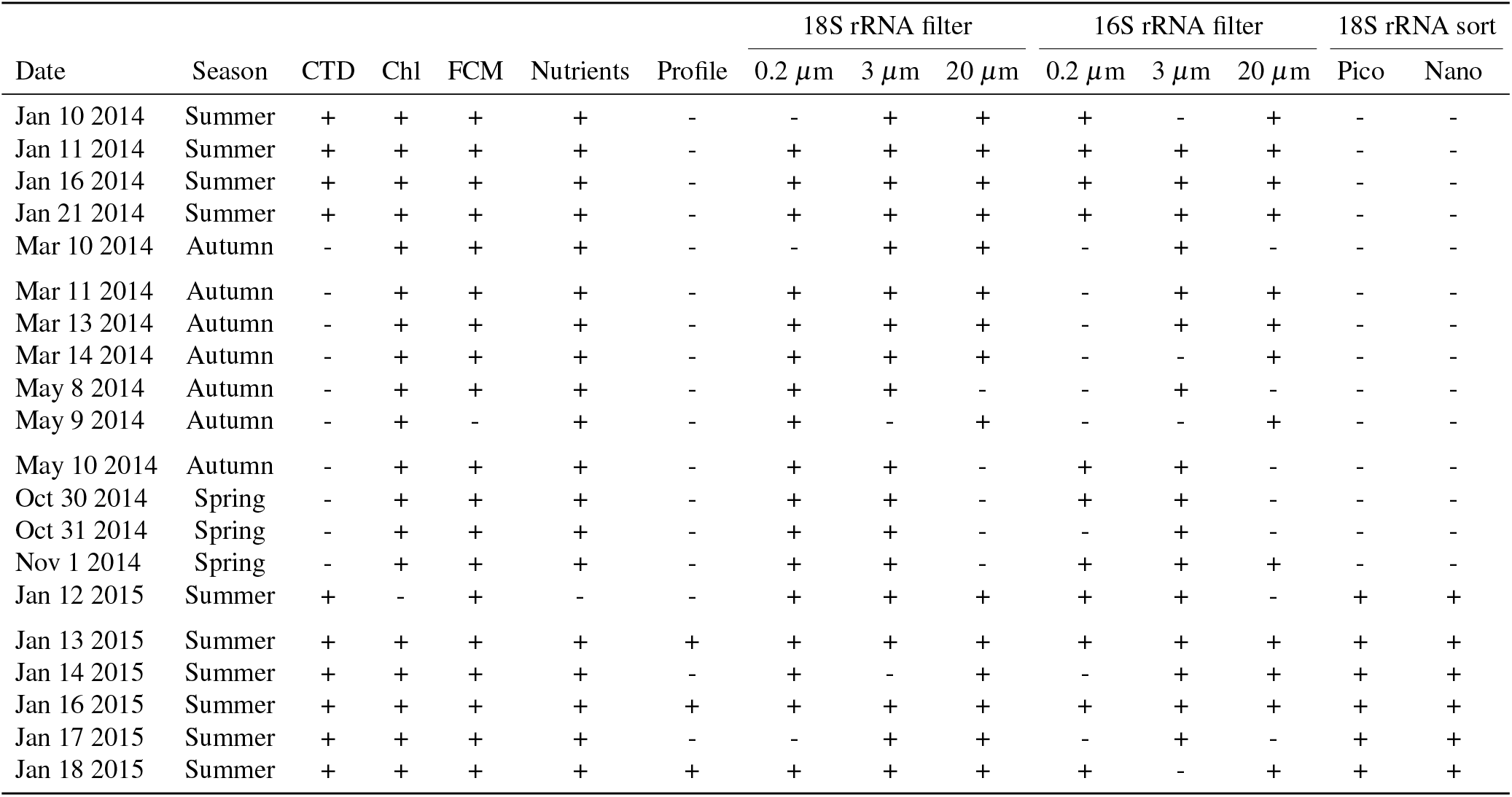
Samples collected. CTD corresponds to salinity and temperature data from CTD cast, Chl to Chlorophyll a, FCM to flow cytometry and Profile to vertical profile sampling. 18S and 16S rRNA columns correspond to metabarcoding analyses for nuclear 18S and plastidial 16S rRNA gene.

| Date | Season | CTD | Chl | FCM | Nutrients | Profile | 18S rRNA filter |  |  | 16S rRNA filter |  |  | 18S rRNA sort |  |
| --- | --- | --- | --- | --- | --- | --- | --- | --- | --- | --- | --- | --- | --- | --- |
| | | | | | | | 0.2 μm | 3 μm | 20 μm | 0.2 μm | 3 μm | 20 μm | Pico | Nano |
| Jan 10 2014 | Summer | + | + | + | + | - | - | + | + | + | - | + | - | - |
| Jan 11 2014 | Summer | + | + | + | + | - | + | + | + | + | + | + | - | - |
| Jan 16 2014 | Summer | + | + | + | + | - | + | + | + | + | + | + | - | - |
| Jan 21 2014 | Summer | + | + | + | + | - | + | + | + | + | + | + | - | - |
| Mar 10 2014 | Autumn | - | + | + | + | - | - | + | + | - | + | - | - | - |
| Mar 11 2014 | Autumn | - | + | + | + | - | + | + | + | - | + | + | - | - |
| Mar 13 2014 | Autumn | - | + | + | + | - | + | + | + | - | + | + | - | - |
| Mar 14 2014 | Autumn | - | + | + | + | - | + | + | + | - | - | + | - | - |
| May 8 2014 | Autumn | - | + | + | + | - | + | + | - | - | + | - | - | - |
| May 9 2014 | Autumn | - | + | - | + | - | + | - | + | - | - | + | - | - |
| May 10 2014 | Autumn | - | + | + | + | - | + | + | - | + | + | - | - | - |
| Oct 30 2014 | Spring | - | + | + | + | - | + | + | - | + | + | - | - | - |
| Oct 31 2014 | Spring | - | + | + | + | - | + | + | - | - | + | - | - | - |
| Nov 1 2014 | Spring | - | + | + | + | - | + | + | - | + | + | + | - | - |
| Jan 12 2015 | Summer | + | - | + | - | - | + | + | + | + | + | - | + | + |
| Jan 13 2015 | Summer | + | + | + | + | + | + | + | + | + | + | + | + | + |
| Jan 14 2015 | Summer | + | + | + | + | - | + | - | + | - | + | + | + | + |
| Jan 16 2015 | Summer | + | + | + | + | + | + | + | + | + | + | + | + | + |
| Jan 17 2015 | Summer | + | + | + | + | - | - | + | + | - | + | - | + | + |
| Jan 18 2015 | Summer | + | + | + | + | + | + | + | + | + | - | + | + | + |

Nutrients (NO_3_^−2^, PO_4_^−3^, SiO_3_^−4^) showed maximum levels during autumn and spring, when lower phytoplankton abundance was recorded, and minimum levels during summer, when phytoplankton abundance was higher (Figure 1C). Silicate was the nutrient with the highest concentration, followed by nitrate and phosphate. Chlorophyll *a* (Chl a) concentration, a proxy of phytoplankton biomass, was below 0.4 mg m^−3^ in autumn and spring. Chl *a* was higher in summer 2014 compared to 2015 (Figure 1D).

### Overall composition of the phytoplankton community

Phytoplankton composition was analyzed by three different metabarcoding approaches (Tables 1 and 2). Filtered samples (3 size fractions) were analyzed using both the nuclear 18S rRNA gene, hereafter 18S-filter, and the plastidial 16S rRNA gene, hereafter 16S-filter, while during summer 2015 we were also able to obtain 18S rRNA sequences from flow cytometry sorted populations (pico- and nano-phytoplankton), hereafter 18S-sort. The sequence data were processed with the dada2 pipeline (Callahan et al. 2016) that cluster reads into amplicon sequence variant (ASV). In this paper, we are focusing on the five major eukaryotic divisions that contain photosynthetic taxa: Ochrophyta (in particular diatoms), Chlorophyta (green algae), Haptophyta, Cryptophyta and Rhodophyta (mostly macroalgae). Because a large fraction of dinoflagellate species are heterotrophic, even within the same genus (Jeong et al. 2010), and Chrysophyceae (Ochrophyta) ASVs were assigned to heterotrophic taxa such as *Paraphysomonas* or *Spumella* and to uncultured clades that are known or hypothesized to be heterotrophic, we have excluded these groups from our analysis. Classes for which all the taxa recovered corresponded to macro-algae were also excluded: Bangiophyceae and Florideophyceae (Rhodophyta), Xanthophyceae and Phaeophyceae (Ochrophyta). The total number of ASVs corresponding to photosynthetic taxa varied from 189 for the sorted samples to 564 for the filtered samples. The average number of reads corresponding to photosynthetic taxa was around 30,000 per sample (Table 2).

**Table 2.** Summary of the metabarcoding data sets analyzed. ID corresponds to the dataset identifier. “Photo ASVs” and “Photo reads” corresponds to the number of ASVs and the mean number of reads assigned to photosynthetic taxa.

| ID | Gene | Sample processing | Fractions | Sample number | Photo ASVs | Photo reads (mean) |
| --- | --- | --- | --- | --- | --- | --- |
| 16 | 18S rRNA nuclear | filtered | 0.2, 3, 20 $\mu\text{m}$ | 120 | 564 | 25494 |
| 17 | 16S rRNA plastidial | filtered | 0.2, 3, 20 $\mu\text{m}$ | 100 | 357 | 34999 |
| 18 | 18S rRNA nuclear | sorted | pico, nano | 40 | 189 | 31787 |

An analysis performed in January 2015 over a vertical profile revealed that the water column was not stratified (Table S1) and that the class composition of the phytoplankton community in each size fraction (Figure S1) was fairly uniform vertically. Therefore surface samples can be considered to be representative of the whole water column. It should be noted however that some species were only found at depth in the euphotic zone samples and not in surface (Table S2).

Phytoplankton communities in WAP coastal waters were highly diverse, with 14 classes and 156 species detected in surface samples (Table S3). The major classes were Bacillariophyta (diatoms), Pelagophyceae and Dictyochophyceae for division Ochrophyta, Mamiellophyceae and Pyramimonadophyceae for division Chlorophyta (green algae), Prymnesiophyceae and Cryptophyceae (Figures 2 and S2).

**Figure 2.**
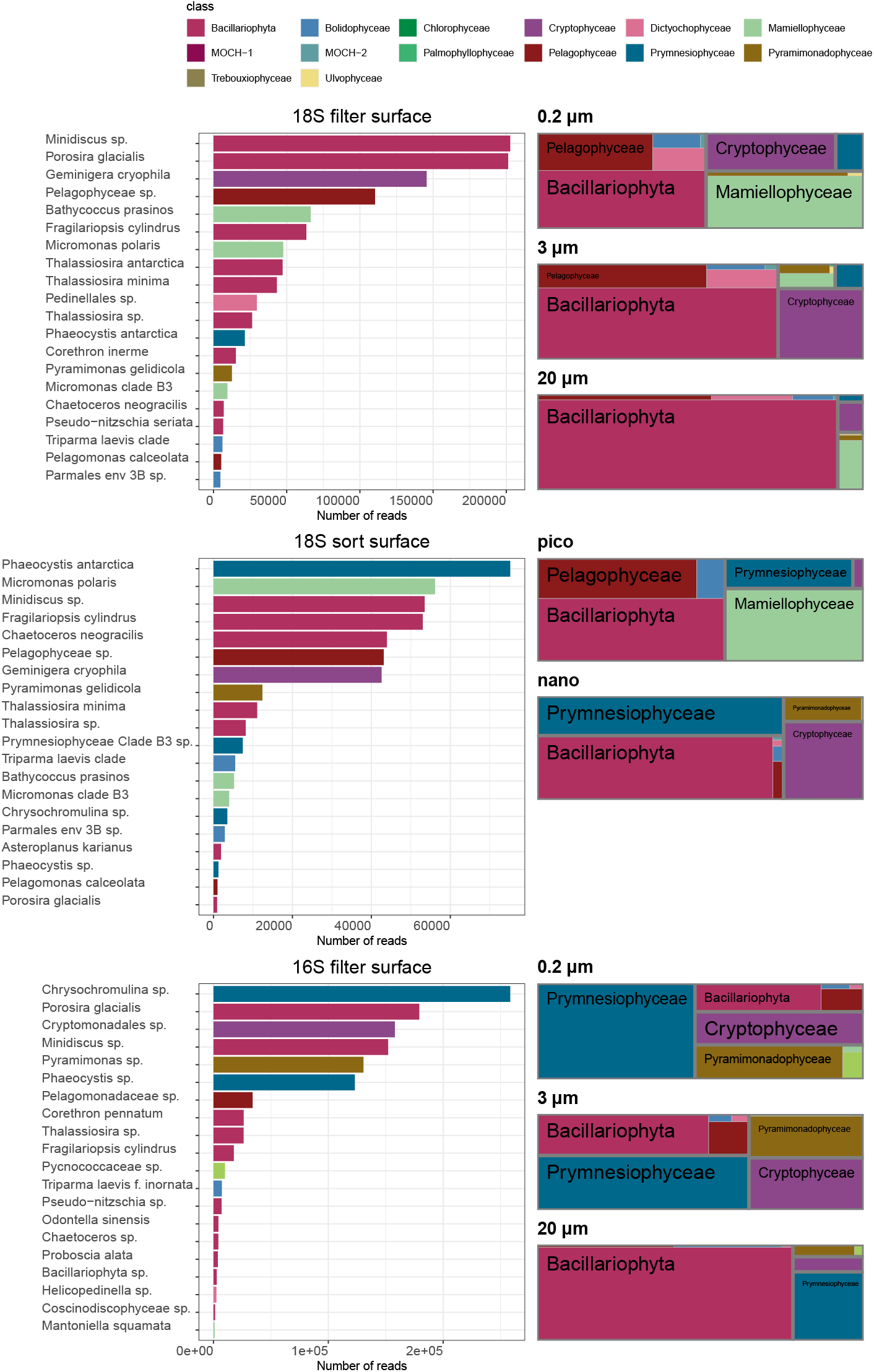
Community composition of phytoplankton at species level for surface samples (5 m) at station 6 in Fieldes Bay. Top panel: 18S rRNA gene for filtered samples. Middle panel: 18S rRNA gene for sorted samples. Bottom panel: plastidial 16S rRNA gene for filtered samples. Left side: abundance rank chart for major species. Right side: proportional area charts of relative abundance of classes by size fraction. 0.2, 3, and 20 *μ*m correspond to the 0.2-3, 3-20 and > 20 *μ*m size fractions, respectively.

Among Ochrophyta, Bacillariophyta were dominating with the species *Porosira glacialis, Fragilar-iopsis cylindrus* and *Chaetoceros neogracilis*, and the genera *Minidiscus* and *Thalassiosira* as major taxa. The sequence of the main ASV assigned to *C. neogracilis* (found in both 18S-filter and 18S-sort datasets) is 100% similar to an Antarctic strain AnM0002 (Genbank EU090012) but differs by 7 positions within the V4 region of 18S rRNA (98.1 % similarity) from all Arctic strains, suggesting that it is a distinct, yet undescribed, species (Figure S3). For some genera such as *Thalassiosira* and *Minidiscus*, the identification down to the species level is difficult because reference sequences are lacking for Antarctic species. The sequence of the main *Minidiscus* ASV (asv_016_00002 from the 18S-filter dataset also found in the 18S-sort dataset) is 100% similar (Figure S4) to strain RCC4582 (Genbank MH843669) which was isolated from Fildes Bay in January 2015. RCC4582 cells are about 5 *μ*m in size and were tentatively identified as *M. chilensis* (unpublished observations). This sequence (asv_016_00002) was also 100% identical to *Shionodiscus oestrupii* var. *venrickiae* strain CC03-15 (Genbank DQ514870) which has larger cells (Wilks and Armand 2017) and therefore is probably mis-identified. Within *Thalassiosira*, the major ASV (asv_016_00006 also present in 18S-sort) is 100% similar to *Thalassiosira antarctica* strain UNC1404 (KX253953) that was isolated off the WAP (Moreno et al. 2018). The second ASV (asv_016_00008 also present in 18S-sort) is 100 % identical to *Thalassiosira minima* strain RCC2265 which was isolated from the Arctic (Balzano et al. 2017) but also to strain RCC4586 which was isolated from Fildes Bay. In contrast, the next *Thalassiosira* ASV (asv_016_00016 also found in 18S-sort) does not match any existing sequence from cultures.

Within Pelagophyceae, two of the major ASV (found in both 18S rRNA datasets) share 99.7 % similarity between them and do not match any described species or even cultured strain, suggesting that they corresponds to a new taxon. One less abundant ASV found in both 18S rRNA datasets matches at 100% *Pelagomonas calceolata*, the type species of this class which is widespread in open oceanic waters (Worden et al. 2012). Among Dictyochophyceae, the main ASV matches with 97.7% similarity *Helicopedinella tricostata* and with higher similarity (99.2%) an undescribed strain (RCC2289) isolated from the Arctic (Balzano et al. 2012a), suggesting that this ASV may correspond to a new species or even genus, while some of the other ASVs match the species *Florenciella parvula* and *Pseudochattonella farcimen.* Bolidophyceae were represented by *Triparma laevis* as well as environmental clades (Kuwata et al. 2018). One uncultivated group MOCH-2 (Marine OCHrophyta, Massana et al. 2014) was found in many filtered and sorted samples although at low abundance.

Among Chlorophyta, Mamiellophyceae dominated with three major taxa: *Micromonas polaris, Micromonas* sp. clade B3 (uncultured) and *Bathycoccus prasinos.* While the main *M. polaris* ASVs (found in both 18S datasets) were 100% identical to Arctic strains, some minor *M. polaris* ASVs have a clearly different signature (Figure S5, arrows). On the other hand, the clade B3 ASVs matched the reference sequences from this clade (Tragin and Vaulot 2019). Among Pyramimonadophyceae, the major ASV (present in both 18S datasets) corresponds to the mixotrophic species *Pyramimonas gelidicola.* The other green classes (Trebouxiophyceae, Chlorophyceae, Ulvophyceae and Palmophyllophyceae) or orders (Pseudoscourfeldiales) were only minor contributors to the community.

*Phaeocystis antarctica* was the dominant Prymnesiophyceae (Haptophyta) species among 18S rRNA metabarcodes (Figure S6). However, in the sorted samples, we also found a minor ASV (asv_018_00239), not present in surface but only at depth (Table S2), with a 100% match to a strain of the arctic species *Phaeocystis pouchetii.* Surprisingly, the sequence of the three dominant Prymnesiophyceae ASVs in the 16S metabarcodes matched *Chrysochromulina throndsenii* with about 98% similarity, while they were matching *P. antarctica* with only 93% similarity. The fourth Prymnesiophyceae ASV (asv_017_00037) matched a *P. antarctica* strain at 100%.

Among Cryptophyceae, the dominant species was *Geminigera cryophila* with small contributions of the genera *Hemiselmis* and *Plagioselmis.* The most abundant ASV (asv_016_00003) found in both 18S-filter and 18S-sort is 99.7% similar to a sequence from a recently isolate of *G. cryophila* from Antarctica (HQ111513, Hoff et al. 2020). Another abundant Cryptophyceae ASV (asv_016_00113) is 100% similar to several strains isolated from the Wedell and Ross Seas, some originating from the ice (e.g. RCC5152). Asv_016_00003 and 00113 were only 98.9% similar. An ASV (asv_017_00002) assigned to Cryptomonadales was also abundant in the 16S dataset, maybe corresponding to *G. cryophila* as well, since it is 99.5 % similar to a sequence from this species (AB073111), although it more similar to *Teleaulax amphioxeia* sequence (99.7 %).

The dominant taxa clearly varied depending on sample processing and the marker gene used (Figure 2, left panels). Filtered samples using the 18S rRNA gene were dominated by the diatoms *Minidiscus* sp., *P. glacialis*, *F. cylindrus*, *T. antarctica* and *T. minima*, the cryptophyte *G. cryophila*, an unknown pelagophyte, and *B. prasinos*. In sorted samples using the 18S rRNA gene, the dominant taxa were *P. antarctica*, followed by *M. polaris, Minidiscus* sp., *F. cylindrus, C. neogracilis* (which was much less abundant in filtered samples) and an unknown pelagophyte. Finally, filtered samples analyzed with 16S rRNA gene were dominated by species from the class Prymnesiophyceae *(Chrysochromulina* sp.) followed by the diatom *P. glacialis*, an unknown cryptophyte and *Minidiscus* sp.

We performed a more detailed analysis at the genus level to compute the number of taxa common to different approaches (Figure 3A). We focused on the summer 2015, the only period for which we have comparable datasets. For the filtered samples, we only considered the 0.2 and 3 *μ*m fractions for comparison with the sorted samples which do not include the microphytoplankton. The number of shared genera detected by the three approaches was low (15, Figure 3A). The number of genera only detected in one approach was highest for the 18S filter dataset (28, in particular diatoms), followed by 16S from filters (8, in particular diatoms and Dictyochophyceae), and 18S from sorted samples (4, three diatoms and one pelagophyte).

**Figure 3.**
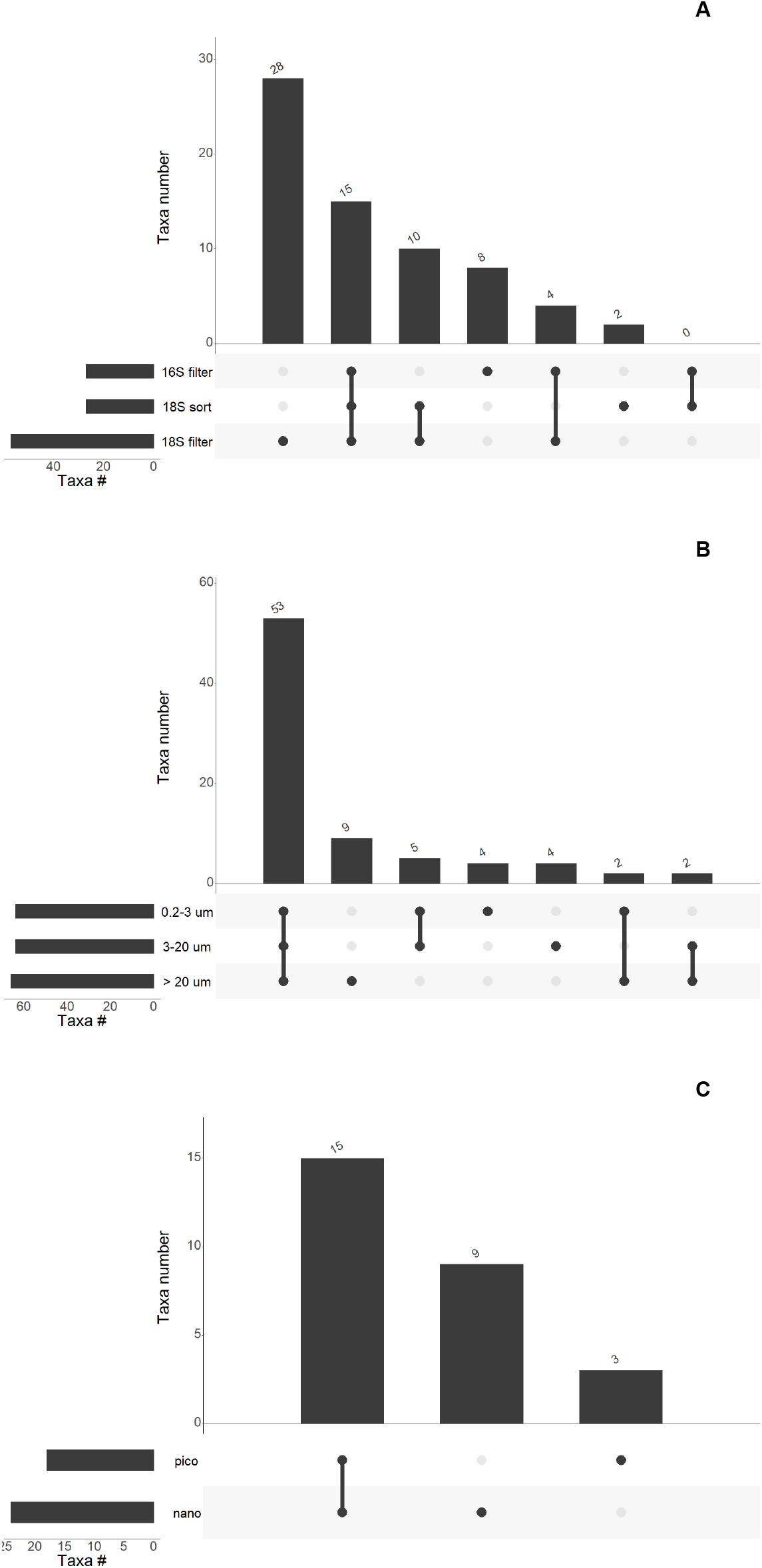
(A) Number of genera in common between different metabarcoding approaches for samples of the 0.2-3 and 3-20 *μ*m size fractions, collected during summer 2015. (B) Number of genera in common between different size-fraction for all 18S rRNA gene samples. (C) Number of genera in common between different populations sorted by flow cytometry in summer 2015. Only taxonomic valid genera have been included.

### Community size structure

In the larger size fractions (20 *μ*m for filtered samples and nano for sorted samples), diatoms were always dominant whatever the metabarcoding approach used (Figures 2 right side, and S2). In the smaller size fraction (0.2 *μ*m and pico), the composition was more dependent on the approach. For example, with both 18S-filter and 18S-sort data, Mamiellophyceae were important but were almost absent in the 16S-filter data. In the filter data, Prymnesiophyceae were much more prevalent with 16S compared with 18S, especially in the two smaller fractions (Figure S2). An analysis of the genera common to different size fractions (Figure 3B) based on 18S reveals that more than 65% of the genera were found in the three size-fractions (53) suggesting that size fractionation is not very efficient at strictly separating phytoplankton communities. When looking at sorted samples (Figure 3C), the same observation prevailed as more than 55% of the genera were found in both pico and nano sorted fractions. This must be tempered however when looking at the abundance of each genus (Figure S2) with many genera abundant only in a single size fraction, although they may present in the other size fractions at low abundance. For example, although *Micromonas* was present in all filtered size fractions and sorted samples (Supplementary Data S2), it was only abundant in the smallest size fractions (Figure S2). Similarly *Porosira* sequences are found in all filtered size fractions (Supplementary Data S2) but dominant in the 20 *μ*m fraction and much lower in the 0.2 *μ*m one.

### Annual dynamics

The dynamics of the phytoplankton community throughout the year could only be followed from the filtered samples since sorted samples were only obtained in summer 2015. The most abundant photosynthetic classes showed a clear seasonal pattern with year to year variation (Figures 4, S7 and S8). Focusing first on the 18S-filter dataset for which we have the largest number of samples, (Figures 4), we observed in the 0.2 *μ*m size-fraction, a succession from Bacillariophyta in summer to Pelagophyceae and Cryptophyceae in the fall and spring, and then back to Bacillariophyta. The main species in this size fraction were *Minidiscus* sp. during summer, an unknown member of the Pelagophyceae during autumn and spring, and *G. cryophyla* during spring. The latter two taxa had also high abundance for the last samples taken in summer 2015. Sequences assigned to Mamiellophyceae were detected throughout all the sampled dates in the 0.2 *μ*m size-fraction. *B. prasinos* was present in the fall and spring. In contrast *M. polaris* was most prevalent during the summer 2015. In the 3 *μ*m fraction, diatoms were only dominant during the summer and early fall while Cryptophyceae were abundant throughout spring 2014 and summer 2015 and Pelagophyceae at the end of the fall and in the spring. In this size fraction, the dominant diatom was *Minidiscus* sp. followed by *F. cylindrus* and *T. minima*, and the dominant cryptophyte *G. cryophyla*. Finally in the 20 *μ*m fraction, diatoms were dominant throughout the year with the exception of the last sample taken in the fall (May 2014) in which pelagophytes peaked. In this fraction, it was the larger diatom *P. glacialis* which was contributing most, followed by *T. antarctica* and the smaller *Minidiscus* sp. Interestingly when looking at the summer, there was some year to year variation. For example, Cryptophyceae were abundant in the summer in 2015 but less so in 2014 while it was the reverse for Dictyochophyceae. The 16S-filter dataset is interesting because while confirming the 18S-filter data, it provides better insight into the seasonal dynamics of Prymnesiophyceae and Pyramimonadophyceae that are masked by other taxonomic groups in the latter dataset (Figure S7). Prymnesiophyceae, especially prevalent in the pico and nano-phytoplankton fractions, are present throughout the year with a peak in the fall while Pyramimonadophyceae, almost absent from the micro-phytoplankton, are restricted to the summer.

**Figure 4.**
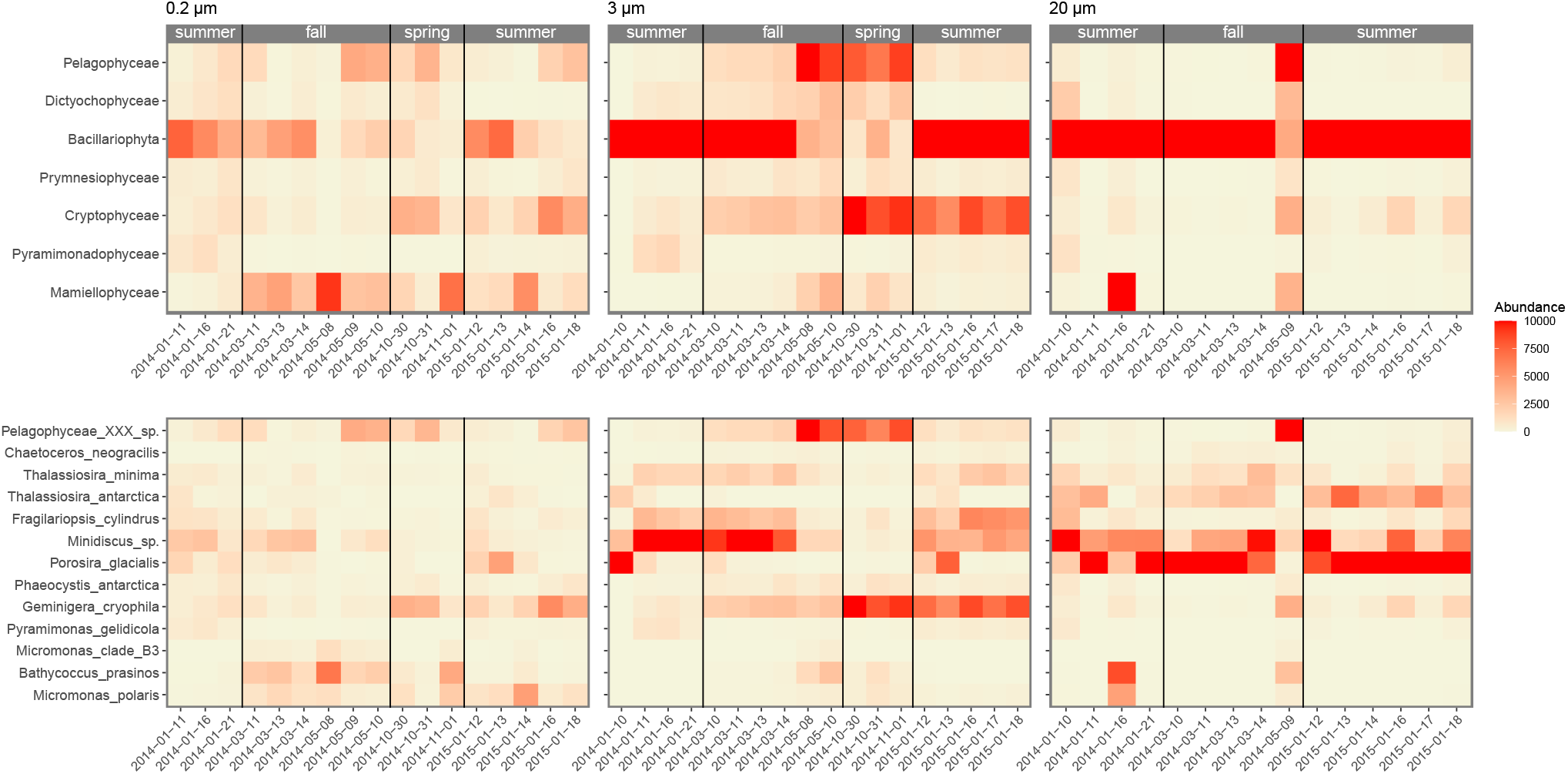
Relative abundance of the main phytoplankton groups at class (top) and genus (bottom) levels in Fildes Bay during the study period. The color scale of the heatmap correspond to the relative abundance of each taxon based on the 18S rRNA gene in filtered surface samples. Left: 0.2-3 μm. Middle: 3-20 μm. Right: > 20 μm. Season delimitation corresponds to meteorological seasons.

NMDS analysis based on Bray-Curtis dissimilarity for 18S-filter metabarcodes (Figure 5 top) shows that samples group together according to season and size fraction with summer samples displaying most scatter. Besides, taxa distribution also shows a seasonal variation, with Bacillariophyta as the dominant class in summer, while Prymnesiophyceae and Cryptophyceae more dominant in the other seasons. When available environmental parameters were fitted against the NMDS analysis, silica and nitrates appear as key factors to differentiate summer vs. spring and autumn. A similar clustering pattern was observed when using the plastidial 16S rRNA gene (Figure S9). Clustering based on either season or size fraction was supported by ANOSIM as highly significant and size fraction had a stronger clustering effect than season (Table S4).

**Figure 5.**
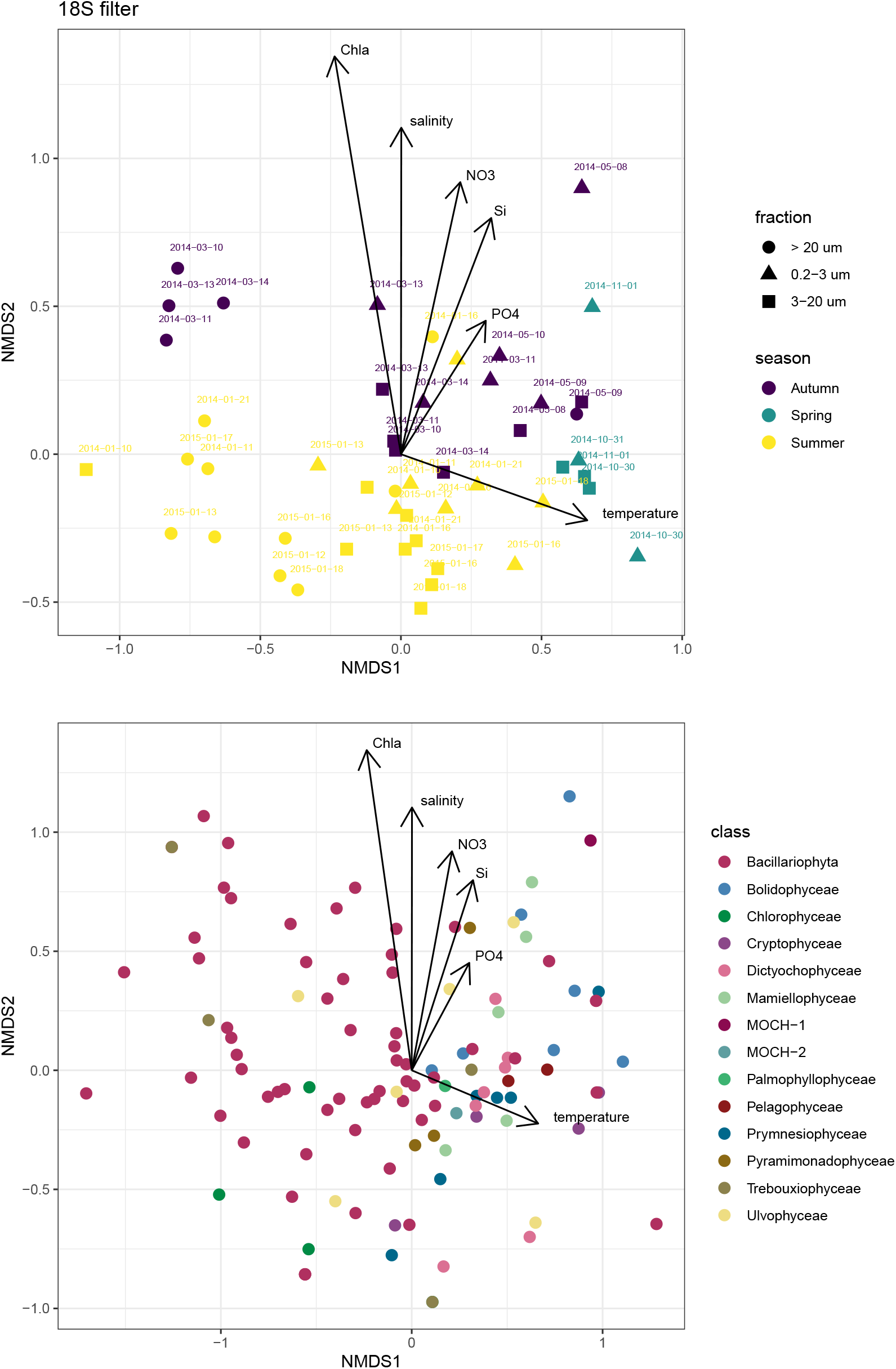
Non-metric multidimensional scaling (NMDS) analysis based on Bray-Curtis dissimilarities of the phytoplankton community composition (species) labeled by meteorological season (summer, autumn, and spring) and size fraction based of the 18S gene of filtered samples. (Top) Samples. (Bottom) ASVs. Stress = 0.16.

## Discussion

In this work, we assessed the variability in phytoplankton abundance, diversity, and community composition during the austral year in a coastal area of the WAP by metabarcoding using two different genes: nuclear 18S rRNA and plastidial 16S rRNA. The community structure determined using these two markers displayed marked differences for some phytoplankton groups like Prymnesiophyceae, Pelagophyceae and Mamiellophyceae. Differences in sequencing results between marker genes have been noted before (Shi et al. 2011), and could be linked to primer bias, differences in amplification efficiency, variations in number of gene copies per genome (Needham and Fuhrman 2016), differences in number of plastid genome copies per cell resulting from differences in the number of chloroplasts par cell (Lin et al. 2019) or differential extraction yield for nuclear vs. plastidial DNA. These differences highlight the interest of using both gene markers for a more complete assessment of phytoplankton community composition. For example, the variation of Prymnesiophyceae and Pyramimonadophyceae over the year was easier to visualize with the 16S-filter dataset, while Mamiellophyceae significant contribution to the phytoplankton community was only evident on the 18S dataset.

These discrepancies point out that the use of different sample processing and marker genes allows to get a more complete image of phytoplankton communities. For example, some groups such as Prymnesiophyceae and Pyramimonadophyceae were more represented when using plastidial 16S versus nuclear 18S while Mamiellophyceae were almost absent from the 16S amplicon data. Pseudoscourfeldiales (Chlorophyta) only appeared in the 16S data. The uncultured marine Ochrophyta (MOCH, Massana et al. 2014), described from environmental 18S rRNA sequences, was also only detected in the 18S data since no 16S rRNA sequences have been attributed to this uncultured clade (Supplementary Data S2).

### Phytoplankton annual succession in Antarctic coastal waters

Phytoplankton composition in the WAP has been studied before (Kopczynska 2008; Lange et al. 2018), but many of these studies relied on optical microscopy and pigment analysis (Biggs et al. 2019; Leeuwe et al. 2020; Rozema et al. 2016; Wasilowska et al. 2015) and focused only on the summer period (Annett et al. 2010; Garibotti et al. 2003; Lima et al. 2019). Metabarcoding characterization in the WAP has been performed for samples from the PAL-TER, Fildes Bay (King George Island) and the RaTS (Luo et al. 2016; Luria et al. 2014; Rozema et al. 2017). However, none of these studies investigated the structure of the phytoplankton community at different seasons. In the present study, succession of different phytoplankton groups through the Austral seasons was evident. Bacillariophyta (diatoms) dominated mainly in summer and early autumn in all fractions; Mamiellophyceae were present in the pico-phytoplankton fraction throughout the year; Pelagophyceae, Dictyochophyceae and, to a lesser extent, Cryptophyceae dominated late autumn and spring samples, while Prymnesiophyceae increased at the end of summer in the small size fraction.

The most abundant genera of diatoms included *Chaetoceros, Thalassiosira, Fragiliaropsis, Minidiscus* and *Porosira*. These genera have been often observed in the WAP during summer months (Annett et al. 2010; Lange et al. 2018), although the exact species may be different. For example, Garibotti et al. (2003) reported that different *Fragilariopsis* species could account together for up to 88% of diatom cell abundance at some sites in WAP during summer. In our study, the main species was *F. cylindrus* while *F. sublineata* was also present but much less abundant (Table S2). We failed to observe other *Fragilariopsis* species often associated to WAP spring/summer blooms, such as *F. pseudonana*, *F. ritscheri* and *F. curta* (Garibotti et al. 2003; Lee et al. 2015). *Minidiscus chilensis* has been previously reported at WAP (Lange et al. 2018) as a characteristic diatom of early-summer production, comprising a high proportion of phytoplankton biomass (Annett et al. 2010) and carbon transport to sea-floor (Kang et al. 2003). However, in contrast to the reported early-summer blooms of *Minidiscus* in Ryder Bay (Annett et al. 2010) and Bransfield Strait (Kang et al. 2003), we detected a high abundance of *Minidiscus* in our summer and early autumn samples.

In the pico-phytoplankton fraction, Mamiellophyceae were present throughout the year and dominated specific samples from autumn and summer, although the most abundant species, *M. polaris* has been rarely reported in Antarctic waters, in contrast to its dominance within the Arctic pico-phytoplankton (see next section). In the pico- and nano-phytoplankton fractions, Pelagophyceae became abundant after diatoms had decreased towards the end of autumn (Figure 4). Pelagophyceae is a class with only a few species described, mostly belonging to the pico-plankton size range (Vaulot et al. 2008), that was initially described from strains isolated in tropical and temperate waters (Andersen et al. 1993). However this class has been found later in polar environments (Balzano et al. 2012b; Diez et al. 2001; Luo et al. 2016) and recently novel nano-plankton sized strains have been isolated from polar waters which probably correspond to several novel species (Balzano et al. 2012a; Gérikas Ribeiro et al. 2020).

Within Prymnesiophyceae, the genus *Phaeocystis* is considered a key-player in Antarctic waters not only during the highly productive summer, but also during autumn and winter months (Sow et al. 2020). *P. antarctica* has a wide presence in the Southern Ocean (Gaebler et al. 2007) and is linked to increased carbon transport to deeper waters (Arrigo et al. 1999; DiTullio et al. 2000). An alternation between diatoms and *P. antarctica*, as reflected here in the 16S-filter prymnesiophytes (Figure S7), has been reported as a consequence to disturbances in the water column structure (Arrigo et al. 2000; Egas et al. 2017), as the latter benefits from deeper mixed layers and weakly stratified waters, due to its ability to maintain its photosynthetic rates in low light environments (Arrigo et al. 1999) and to quickly acclimate to different light regimes even under iron limitation (Van Leeuwe and Stefels 2007). The shift of prymnesiophytes from the 3 to the 20 *μ*m size fraction in the early summer and late fall 2014 (Figure S7) could be due to the formation of *Phaeocystis* colonies of large size that were retained by the 20 *μ*m filter. Differences observed between genomic 18S rRNA and plastidial 16S rRNA *Phaeocystis* read abundance might be a result of this photo-acclimation process, as an increased number of chloroplasts will result in an increased 16S/18S rRNA ratio (Lin et al. 2019).

As light availability decreases towards autumn/winter, mixotrophy becomes a possible strategy for photosynthetic organisms to survive during the long period of darkness. Few studies however have been performed on this process (Gast et al. 2014). In the present study, three groups have been reported as possessing mixotrophic species: cryptophytes, dictyochophytes and Pyramimonadophyceae. Cryptophyte blooms are considered a secondary stage of the seasonal phytoplankton succession, developing after sea-ice edge diatom blooms, and may present a significant inter-annual variability at WAP, being favored by years of earlier sea-ice retreat (Garibotti et al. 2005). Our data are coherent with this pattern as cryptophytes were most abundant in the spring, when the sea-ice melts. Interestingly, they remained abundant in the summer of 2015 but not in 2014, pointing to some inter-annual variability. *G. cryophila* was the main cryptophyte species in this study, and has been determined to be mixotrophic (Gast et al. 2014). It has been previously reported at WAP (Egas et al. 2017), including as a dominant taxa (Luo et al. 2016), and has probably a circum-Antarctic distribution (Hoff et al. 2020), linked to warmer, nutrient-depleted post-bloom conditions (Gast et al. 2014). Dictyochophyceae were most abundant in the spring under low light conditions. Some of the main ASVs were assigned to Pedinellales, which are known mixotrophs (Sekiguchi et al. 2003) and also to the genus *Florenciella*, which have been very recently determined to be mixotrophic feeding on bacteria as well as cyanobacteria (Li et al. 2020). In contrast, Pyramimonadophyceae which harbor several mixotrophic species (Gast et al. 2014; Maruyama and Kim 2013) were most abundant in the summer, suggesting that the occurring species were probably not mixotrophic.

### Antarctic vs. Arctic phytoplankton communities

The Arctic and Antarctic marine ecosystems share many similarities due to the constraints of solar radiation input at high latitudes and a phytoplankton phenology connected to sea-ice formation and melting. This similarity is also seen at the taxonomic level, as many of the dominant taxa observed in the present study shared highly related or identical 18S rRNA sequences to Arctic species. Bipolarity has been long observed on planktonic marine organisms (Darling et al. 2000; Sul et al. 2013), and implies trans-equatorial genetic flow and organismal dispersal, mainly via ocean currents. Bipolar species might however thrive differently in the Arctic and Antarctic. In a study investigating bipolar protists based on 18S rRNA, Wolf et al. (2015) observed that only two OTUs that were not part of the rare biosphere, i.e. that accounted for more than 1% of total reads, were found in both poles: an unknown alveolate and *Micromonas*.

Although the dominant component of the picophytoplankton in Arctic waters in summer (Balzano et al. 2012b; Lovejoy and Potvin 2011), *M. polaris* has been rarely reported from Antarctic waters (Delmont et al. 2015; Simmons et al. 2015), and even then, in low abundance (Luo et al. 2016; Rozema et al. 2016). In the present study *M. polaris* was detected in 42 samples, reaching up to 47% of photosynthetic reads in a single sample (Table S3). Two other *Micromonas* clades have been detected in Arctic or sub-Arctic waters, clade B3 (Tragin and Vaulot 2019), also detected here, and *M. commoda* clade A2 (Joli et al. 2017). To the best of our knowledge, this is the first study to report this genus as a major player within the austral pico-phytoplankton. It is unclear if the unprecedented high abundance of *M. polaris* in Antarctic waters is related to a local and transient phenomena or part of a greater change associated with global climate patterns, since this species seems to be favored by increasing temperatures, enhanced water column stratification and ocean acidification (Benner et al. 2019; Hoppe et al. 2018; Li et al. 2009). We have also detected a third *Micromonas* signature, which could potentially represent a novel Antarctic *Micromonas* clade (Figure S5). Another Mamiellophyceae, *B. prasinos*, is widely distributed in the world’s oceans with two ecotypes reported so far which share identical 18S rRNA sequences but differ in their genomes and distribution (Vannier et al. 2016; Vaulot et al. 2012). In the present study, *B. prasinos* was abundant during autumn and spring, whereas *M. polaris* was more abundant during spring and summer. Interestingly, *Micromonas* clade B3 seems to follow seasonal dynamics that are closer to *B. prasinos* than to *M. polaris.* These seasonal dynamics seem to be analogous to what was observed in the Arctic, where a seasonal succession occurs between the two taxa with increased abundance of the *Bathycoccus* in winter (Joli et al. 2017), possibly due to differences in loss rates, viral defense efficiency or mixotrophic activity between the two species.

The large centric diatom *Porosira glacialis*, which has a bipolar distribution, was the most abundant taxon in the present data set, mainly in the 20 *μ*m size fraction, reaching up to 74% of total reads in a given sample (Table S3). In the Arctic, *P. glacialis* has been reported as highly abundant in spring samples, co-occurring with *Thalassiosira gravida/antarctica var. borealis* (Kauko et al. 2018). A similar trend was observed in Antarctica, where *P. glacialis* was reported along with *T. antarctica* to make up to 90% of total phytoplankton biomass on King George Island during episodic events (Schloss et al. 2014). These diatoms are considered summer/autumn bloom species which share similar ecological preferences, being found together in diatom assemblages from paleontological samples (Świło et al. 2016). The alternation between *P. glacialis* and *T. antarctica* dominance seems to be linked to sea-ice concentration, as *P. glacialis* higher abundances are correlated to cooler environmental conditions (Pike et al. 2009). Although being often reported from both poles, Arctic and Antarctic strains of *P. glacialis* might differ in their 28S rRNA sequence, indicating a possible genetic divergence (Balzano et al. 2017).

*C. neogracilis* is a species complex with identical 18S rRNA gene, common in Arctic surface waters in the summer (Balzano et al. 2012b, 2017; Lovejoy and Potvin 2011). The *C. neogracilis* partial 18S rRNA sequence obtained in the present study is identical to a previously isolated *C. neogracilis* Antarctic strain (AnM0002), which is morphologically similar to, but phylogenetically distinct from, Arctic strains. Balzano *et al.* (2017) sequenced the full 18S rRNA gene of the AnM0002 strain and reported a 98.9% sequence identity with Arctic *C. neogracilis* strains, suggesting the former could be an undescribed *Chaetoceros* species, possibly with an endemic Antarctic distribution.

*Thalassiosira* spp. is a well-known and important component of both Arctic (Luddington et al. 2016) and Antarctic (Kopczynska 2008; Lange et al. 2018) phytoplankton communities. In the present study *T. minima* was the most conspicuous species among the genus *Thalassiosira*, observed in 49 samples (Table S3). *T. minima* is considered a cosmopolitan species mostly observed in temperate waters (Hoppenrath et al. 2007; Luddington et al. 2016) and mostly excluded from polar regions except for one report in the Arctic Beaufort Sea (Balzano et al. 2017). Surprisingly, *T. minima* does not seem to have been reported in the Southern Ocean which could point out to a recent invasion linked to global change.

*Phaeocystis* is an ubiquitous genus, with a relatively well-defined biogeographic distribution for some species (Schoemann et al. 2005). *P. pouchetii* is mainly found in Arctic and *P. antarctica* in many regions of the Southern Ocean (Gaebler et al. 2007; Lange et al. 2002; Schoemann et al. 2005), while *P. globosa* is mostly found in temperate and tropical waters (Medlin et al. 1994). Although the main ASVs found in this study matched *P. antarctica* confirming many previous reports, we also found one ASV matching *P pouchetii*, the Arctic species, and which was only found at depth (Table S2) suggesting that this latter species might be bipolar.

### Final considerations

The WAP is undergoing accelerate environmental changes compared to the rest of Antarctic regions, being more susceptible to warming and sea-ice loss (Thompson and Solomon 2002) due to increased maritime influence (Smith and Stammerjohn 2001). The decreasing sea-ice extent in both time and space influences phytoplankton diversity and production (Rozema et al. 2016), highlighting the need for year-round ecological assessments of the phytoplankton structure and possible climate-related disturbances. The present study provides evidence that groups such as Mamiellophyceae and Pelagophyceae may have a greater ecological importance in the WAP than previously thought, and that a combination of methods are needed to investigate the full extent of phytoplankton diversity in this region.

## Methods

### Study site and sampling

Surface seawater samples (5 m) were collected from Fildes Bay, King George Island, Western Antarctic Peninsula (62 ° 12’11”S, 58 °55’15”W) using a 5 L Niskin bottle, in January, March, May, and October 2014, and January 2015 (Table 1). In January 2015, vertical profiles were also obtained by sampling at 4 additional depths (15, 20, 25 and 50 m). Samples were prefiltered on board using a 100 *μ*m Nitex mesh, stored in sterile plastic carboys and kept in darkness until processing (less than 2 hours). Once in the laboratory, sub-samples for Chl a, flow cytometry, nutrients and molecular analyses were taken. Temperature (SST), salinity and PAR measurements were obtained using a CTD SBE 911 plus (SeaBird Electronics) equipped with an auxiliary biospherical PAR sensor.

### Nutrients

Sub-samples of filtered seawater were collected in 15 mL polypropylene tubes and stored at −20 °C until analysis. Concentrations of nitrate NO^−3^, phosphate PO_4_ ^−3^ and silicate SiO_2_ ^−4^ were determined as described previously (Hansen et al. 2012).

### Chlorophyll *a* determination

Total Chl *a* was determined from triplicate 100 mL sub-samples. Biomass (<100 μm) was collected on 25 mm diameter GF/F filters (Whatman) in the dark immediately after the samples arrived to the laboratory. Pigments were extracted in 90% acetone for 24 h at −20 °C and analysed on a Turner Designs Trilogy fluorometer, according to the method of Holm-Hansen *et al.* (1965). Calibration was made with a Chl *a* standard (Sigma-Aldrich).

### Phytoplankton cell counts by flow cytometry

Sub-samples of 1.35 mL were taken in triplicates, fixed with 150 *μ*l of fixative (final concentrations: 1% paraformaldehyde, 0.5% glutaraldehyde, 100 mM sodium borate, pH 8.4), incubated for 20 min at room temperature and fast frozen in liquid nitrogen. Cells were enumerated using an Accuri C6 Plus flow cytometer (Becton Dickinson) equipped with a combination of blue 488 nm and red 640 nm lasers. Photosynthetic pico-eukaryotes (PPE), photosynthetic nano-eukaryotes (PNE) and cryptophytes (CRY) were differentiated by forward and side light scatters and trigger pulse width from the 488 nm laser, and red (>670 nm) and orange (585/40 nm) fluorescence detection from 488 and 640 nm laser. Each sample was run at an average flow rate of 81 *μ*L min^−1^ for 3 min. Flow rate was calculated by measuring the difference of volume of pre-filtered water after run for 10 minutes at the fast flow speed. Cell count analyses were performed using BD CSampler Plus software.

### Sorting by flow cytometry

Samples (1.5 mL) for cell sorting by flow cytometry were collected in cryotubes with 10% DMSO (final concentration) and 0.01 % Pluronic F68 (final concentration) (Marie et al. 2014), flash-frozen in liquid nitrogen, and stored at −80°C until analysis at the Station Biologique de Roscoff, France. Samples were analyzed and sorted using a FACSAria flow cytometer (Becton Dickinson, San Jose, CA). Photosynthetic pico and nanoeukaryotes populations were selected based on light scatter, orange phycoerythrin, and red chlorophyll fluorescence and sorted in purity mode, directly into Eppendorf tubes containing Tris-EDTA lysis buffer (Tris 10 mM, EDTA 1 mM, and 1.2% Triton, final concentration). Tris–HCl 50 mM, pH 8.0, NaCl 10 mM was used as sheath liquid. Sheath pressure was set at 70 PSI and nozzle frequency was 90 KHz with a deflection voltage of 6,000 V. Sheath fluid samples were collected and analyzed as negative controls in all subsequent steps, including sequencing, to test for contamination in the flow sorting process (2018).

### Biomass collection and DNA extraction

Samples of 4.5 L of seawater were serially size-fractionated using a peristaltic pump (Cole-Palmer) with 47 mm diameter Swinnex filter holder (Millipore), and 20 *μ*m (Nylon, Millipore), 3 *μ*m and 0.2 *μ*m (Poly-carbonate, Millipore) pore size filters. Filters were stored in 2 mL cryovials in liquid nitrogen or at −80°C until DNA extraction. For DNA extraction, filters were thawed and half of the filters were cut into small pieces, while the other half was kept at −20 °C as backup. All steps were performed under sterile conditions. Each half-filter was incubated in lysis buffer (TE 1x / NaCl 0.15 M), with 10% SDS and 20 mg mL^−1^ proteinase K at 37 °C for 1 h. DNA was extracted using 5 M NaCl and hexadecyl-trimethyl-ammonium bromide (CTAB) extraction buffer (10% CTAB, 0.7% NaCl) and incubated at 65 °C for 10 min before protein removal using a conventional phenol-chloroform method. DNA was precipitated using ethanol at −20 °C for 1 h and re-suspended in 50 *μ*l Milli-Q water (Millipore) (Egas et al. 2017). DNA integrity was evaluated by agarose gel electrophoresis and quantified using a fluorometric assay (Qubit 2.0 fluorometer).

### Metabarcoding of filtered samples

For general eukaryotes, the V4 region of 18S rRNA gene was amplified using primer pair TAReuk454FWD1 (CCAGCASCYGCGGTAATTCC) and V4 18S Next.Rev (ACTTTCGTTCTTGATYRATGA) (Piredda et al. 2017). For photosynthetic eukaryotes, plastidial 16S rRNA gene was amplified using primer pair Pla491F (GAGGAATAAGCATCGGCTAA, (Fuller et al. 2006)) and PP936R (CCTTTGAGTTTCAYYCTTGC) (https://biomarks.eu/pp936r). PCR reactions were performed in triplicate in 50 *μ*L final volumes with Taq buffer 1X final concentration, 2 mM of MgCl_2_, 0.2 nM of dNTPs, 0.2 *μ*M of each primer, 2.5 units of GoTaq Flexi DNA Polymerase (Fermelo) and approximately 5 ng μl^−1^ of DNA. Amplification conditions were 10 min of initial denaturation at 94 °C, 30 cycles of 94 °C for 45 s, 57 °C (V4 18S rRNA) or 62 °C (16S rRNA) for 45 s and 72 °C for 1.25 min, followed by a final extension of 72 °C for 10 min. Amplicons were visualized on a 2% agarose gel (TAE 1X) and purified using the Wizard SV Gel and PCR Clean-Up System.

### Metabarcoding of sorted samples

DNA from sorted cells was extracted by one cycle of freezing and thawing in liquid nitrogen a prior the PCR reaction. Because of the small number of cells collected (from to 500 to 6,500), sorted samples required a nested amplification with the first round of PCR done using the 18S rRNA gene primers 63F (ACGCTTGTCTCAAAGATTA) and 1818R (ACGGAAACCTTGTTACGA) (Lepère et al. 2011) with the following 10 *μ*L mix: 5 *μ*L KAPA HiFi HotStart ReadyMix 2x, 0.3 *μ*M final concentration of each primer, 1 *μ*L of cells. Thermal conditions were: 95 °C for 5 min, followed by 25 cycles of 98 °C for 20 s, 52 °C for 30 s, 72 °C for 90 s, and a final cycle of 72 °C for 5 min. For the second round: 12.5 *μ*L KAPA HiFi HotStart ReadyMix 2x, 0.3 *μ*M final concentration of the same primers as described above (TAReuk454FWD1 and V4 18S Next.Rev), 2.5 *μ*L of first round product and water for a 25 *μ*L reaction. Thermal conditions were: 95 °C for 3 min, followed by 25 cycles of 98 °C for 20 s, 65 °C for 1 min, 72 ^°^C.

### Amplicon sequencing

Amplicons were sequenced on an Illumina Miseq using a 250 cycles Miseq kit v.2 at the Genotoul GeT core facility (Toulouse, France) for filtered samples and at the GenoMer platform (Roscoff, France) for sorted samples. The final amplicon sequencing dataset (Table 2) contained 120 filtered samples (data set # 16) and 40 sorted samples for the 18S rRNA gene (data set # 18), and 100 filtered for the plastidial 16S rRNA gene (data set # 17). See Supplementary Data S1 for list of samples. Data have been deposited to GenBank SRA under project numbers PRJNA645244 for 18S rRNA and PRJNA645261 for 16S rRNA.

### Sequence processing

The three different datasets (16, 17 and 18) were processed independently. Primer sequences were first removed using cutadapt (Martin 2011). Amplicon processing was performed under the R software (R Development Core Team 2013) using the dada2 package (Callahan et al. 2016) with the following parameters: truncLen and minLen = c(230, 230), truncQ = 2, maxEE = c(10, 10). Taxonomic assignation of ASVs was performed using the assignTaxonomy function from dada2 against the PR_2_ database (Guillou et al. 2013) version 4.12 (https://pr2-database.org/) which contains both 18S rRNA and plastidial 16S rRNA reference sequences, the latter originating from a curated version of Phytoref (Decelle et al. 2015). We selected only ASVs corresponding to photosynthetic groups (divisions Chlorophyta, Cryptophyta, Rhodophyta, Haptophyta and Ochrophyta with the exception of Chrysophyceae, Bangiophyceae, Florideophyceae, Xanthophyceae and Phaeophyceae that are known to be either heterotrophic or only contain macroalgae). The number of photosynthetic ASVs and the average number of reads per dataset is provided in Table 2.

### Data analysis

The following R packages were used for data analysis: *tidyr* for filtering and plotting, *treemapify* for treemaps, *phyloseq* (McMurdie and Holmes 2013) for heatmaps and NMDS, *vegan* for ANOSIM (ANalysis Of SIMilarity) and *upsetR* for upset plots. Samples were first normalized by the median sequencing depth.

## Acknowledgements

This work was funded by INACH RG_31-15 and INACH RT_34-17 grants. Collaboration between Chile and France was funded through ECOS-CONICYT No. C16B02 and CNRS International Research Network “Diversity, Evolution and Biotechnology of Marine Algae” (GDRI No. 0803). The authors thank Dr. Ernesto Molina for sampling during autumn and spring, as well the logistic support at the scientific station Professor Julio Escudero, INACH. Adriana Lopes dos Santos was supported by the Singapore Ministry of Education, Academic Research Fund Tier 1 (RG26/19). We thank the Roscoff ABIMS platform of the FR2424 (CNRS, Sorbonne Université) for bioinformatics resources.

## Author contributions statement

NT, RDI, ALS and DV designed the study. MM, RDI, ALS, DV and NT collected the samples. AC and DM performed the flow cytometry analysis. MM, RDI, ALS, RDI, CG, DV and NT performed the data analysis and interpretation. NT, RDI, ALS, CG and DV wrote the paper. All authors read and approved the final manuscript.

## Additional information

**Accession codes**: GenBank project numbers PRJNA645244 and PRJNA645261.

**Code and processed data**: https://github.com/vaulot/Paper-Trefault-2020-Antarctica

**Competing interests** The authors declare no competing financial interests.

## List of Tables

Table. 1 Samples collected. CTD corresponds to salinity and temperature data from CTD cast, Chl to Chlorophyll *a*, FCM to flow cytometry and Profile to vertical profile sampling. 18S and 16S rRNA columns correspond to metabarcoding analyses for nuclear 18S and plastidial 16S rRNA gene.

Table. 2 Summary of the metabarcoding data sets analyzed. ID corresponds to the dataset identifier. “Photo ASVs” and “Photo reads” corresponds to the number of ASVs and the mean number of reads assigned to photosynthetic taxa.

Table. S1 Metadata available for the vertical profile samples of January 16, 2015. PPE, PNE, CRY corresponds to abundance of photosynthetic pico-eukaryotes, nano-eukaryotes and cryptophytes, respectively, in cell mL^−1^.

Table. S2 List of species in the metabarcoding data sets only found in the deep samples (from 10 to 50 m).

Table. S3 List of species found in the metabarcoding data sets for the surface samples. Minimum (min), mean (mean) and Maximum (max) contribution (in %) to the photosynthetic metabarcodes and the number of samples (n) where found for the 18S-filter, 16S-filterand 18S-sort datasets.

Table. S4 ANOSIM analysis for surface samples contrasting the effect of season or size-fraction.

## List of Figures

Fig. 1 Location of the sampling station in Fildes Bay, King George Island, Western Antarctic Peninsula (WAP) and biotic and abiotic characteristics between January 2014 and January 2015. (A) Map of the Antarctica Peninsula and location of the station in Fildes Bay sampled in this study. (B) Phytoplankton abundance measured by flow cytometry. Detected populations correspond to PPE = photosynthetic picoeukaryotes, PNE = photosynthetic nano-eukaryotes, and CRY = cryptophytes. (C) *Nutrients (silicate, SiO_3_^−2^; nitrate, NO_3_ and phosphate, PO_4_^−3^). (D) Chlorophyll a* levels during the sampling period. Values correspond to < 100 *μ*m biomass. For B, C, and D, values represent mean ± standard deviation.

Fig. 2 Community composition of phytoplankton at species level for surface samples (5 m) at station 6 in Fieldes Bay. Top panel: 18S rRNA gene for filtered samples. Middle panel: 18S rRNA gene for sorted samples. Bottom panel: plastidial 16S rRNA gene for filtered samples. Left side: abundance rank chart for major species. Right side: proportional area charts of relative abundance of classes by size fraction. 0.2, 3, and 20 *μ*m correspond to the 0.2-3, 3-20 and > 20 *μ*m size fractions, respectively.

Fig. 3 (A) Number of genera in common between different metabarcoding approaches for samples of the 0.2-3 and 3-20 *μ*m size fractions, collected during summer 2015. (B) Number of genera in common between different size-fraction for all 18S rRNA gene samples. (C) Number of genera in common between different populations sorted by flow cytometry in summer 2015. Only taxonomic valid genera have been included.

Fig. 4 Relative abundance of the main phytoplankton groups at class (top) and genus (bottom) levels in Fildes Bay during the study period. The color scale of the heatmap correspond to the relative abundance of each taxon based on the 18S rRNA gene in filtered surface samples. Left: 0.2-3 μm. Middle: 3-20 μm. Right: > 20 μm. Season delimitation corresponds to meteorological seasons.

Fig. 5 Non-metric multidimensional scaling (NMDS) analysis based on Bray-Curtis dissimilarities of the phytoplankton community composition (species) labeled by meteorological season (summer, autumn, and spring) and size fraction based of the 18S gene of filtered samples. (Top) Samples. (Bottom) ASVs. Stress = 0.16.

Fig. S1 Community composition of phytoplankton at the class level along a vertical profile obtained on January 16, 2015, from 5 m and down to 50 m, based on the 18S rRNA gene for filtered samples.

Fig. S2 Relative abundance of the different genera in surface samples based on three different amplicon sequencing approaches for each size fraction. Left: 18S rRNA gene on filtered samples. Middle: 18S rRNA gene on sorted samples. Right: plastidial 16S rRNA gene on filtered samples.

Fig. S3 Sequence alignment of 18S rRNA ASVs for *Chaetoceros neogracilis* showing the differences between Arctic and Antarctic strains sequences. The ASVs from this study are identical to the Antarctic strain and show 7 bp differences to Arctic strains.

Fig. S4 Sequence alignment of 18S rRNA ASVs for major *Thalassiosira* and *Minidiscus* ASVs in comparison to reference sequences.

Fig. S5 Sequence alignment of 18S rRNA ASVs for *Micromonas* showing the clear signatures for *M. polaris* and clade B3 (Tragin and Vaulot 2019). Within *M. polaris* some sequences have a different signature pointing to a new clade specific of Antarctic waters (arrow).

Fig. S6 Sequence alignment of 18S rRNA ASVs for *Phaeocystis* showing the clear signatures for *P. antarctica* and *P. pouchetii*.

Fig. S7 Read numbers for the main photosynthetic taxonomic groups at the class level for plastidial 16S rRNA gene for filtered surface samples. The color scale of the heatmap corresponds to the normalized number of reads of each taxon. Season delimitation corresponds to meteorological seasons. Left: 0.2-3 μm. Middle: 3-20 μm. Right: > 20 μm.

Fig. S8 Read numbers for the main photosynthetic taxonomic groups at the class (Top) and genus (Bottom) levels of 18S rRNA gene for sorted samples from surface waters. The color scale of the heatmap corresponds to the normalized number of reads of each taxon. Left: pico size fraction. Right: nano size fraction.

Fig. S9 Non-metric multidimensional scaling (NMDS) analysis based on Bray-Curtis dissimilarities of the phytoplankton community composition (species) labeled by meteorological season and size fraction using the plastidial 16S rRNA gene. (A) Samples. (B) ASVs. Stress = 0.15.

## Supplementary material

### Supplementary Data

All supplementary material is available at https://github.com/vaulot/Paper-Trefault-2020-Antarctica

1. **Supplementary Data S1**: List of metabarcoding samples with environmental data (Antarctica_2015_samples.xlsx).
2. **Supplementary Data S2**: List of classes, genera and species found by each metabarcoding approach in surface samples from summer 2015. Taxa with uncertain affiliation (labelled by _X in the PR2 database) were not taken into account (dada2/method_comparison.xlsx).
3. **Supplementary Data S3**: List of ASVs for 18S rRNA gene of filtered samples with abundance table for the different samples - see for sample codes in Supplementary Data S1 (dada2/metapr2_wide_asv_set_16_photo.xlsx).
4. **Supplementary Data S4**: List of ASVs for plastidial 16S rRNA gene of filtered samples with abundance table for the different samples - see for sample codes in Supplementary Data S1 (dada2/metapr2_wide_asv_set_17_photo.xlsx).
5. **Supplementary Data S5**: List of ASVs for plastidial 16S rRNA gene of filtered samples with abundance table for the different samples - see for sample codes in Supplementary Data S1 (dada2/metapr2_wide_asv_set_18_photo.xlsx).
6. **Supplementary Data S6**: Script used to process the data with output (R markdown): https://vaulot.github.io/Paper-Trefault-2020-Antarctica/Antarctica-phyloseq.html.

**Table S1.**
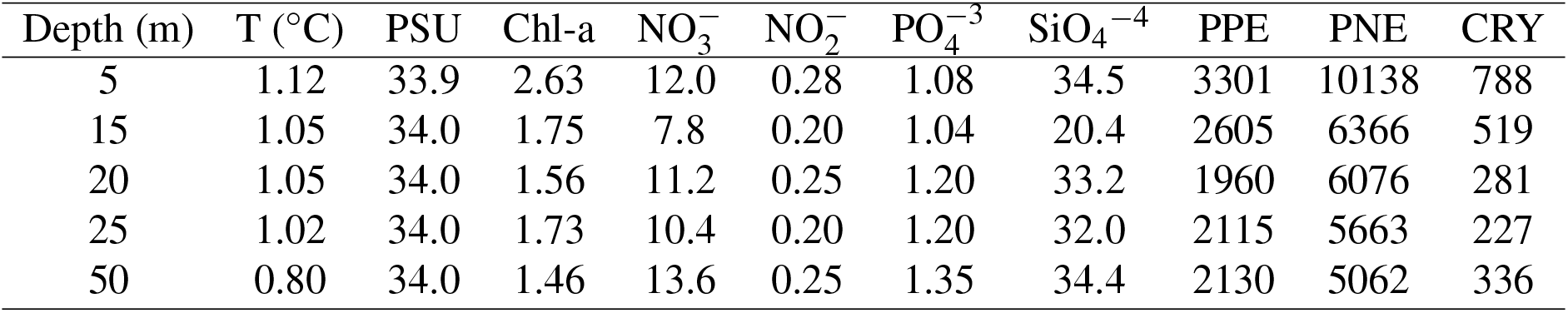
Metadata available for the vertical profile samples of January 16, 2015. PPE, PNE, CRY corresponds to abundance of photosynthetic pico-eukaryotes, nano-eukaryotes and cryptophytes, respectively, in cell mL^−1^.

| Depth (m) | T (°C) | PSU | Chl-a | NO<sub>3</sub><sup>-</sup> | NO<sub>2</sub><sup>-</sup> | PO<sub>4</sub><sup>-3</sup> | SiO<sub>4</sub><sup>-4</sup> | PPE | PNE | CRY |
| --- | --- | --- | --- | --- | --- | --- | --- | --- | --- | --- |
| 5 | 1.12 | 33.9 | 2.63 | 12.0 | 0.28 | 1.08 | 34.5 | 3301 | 10138 | 788 |
| 15 | 1.05 | 34.0 | 1.75 | 7.8 | 0.20 | 1.04 | 20.4 | 2605 | 6366 | 519 |
| 20 | 1.05 | 34.0 | 1.56 | 11.2 | 0.25 | 1.20 | 33.2 | 1960 | 6076 | 281 |
| 25 | 1.02 | 34.0 | 1.73 | 10.4 | 0.20 | 1.20 | 32.0 | 2115 | 5663 | 227 |
| 50 | 0.80 | 34.0 | 1.46 | 13.6 | 0.25 | 1.35 | 34.4 | 2130 | 5062 | 336 |

**Table S2.**
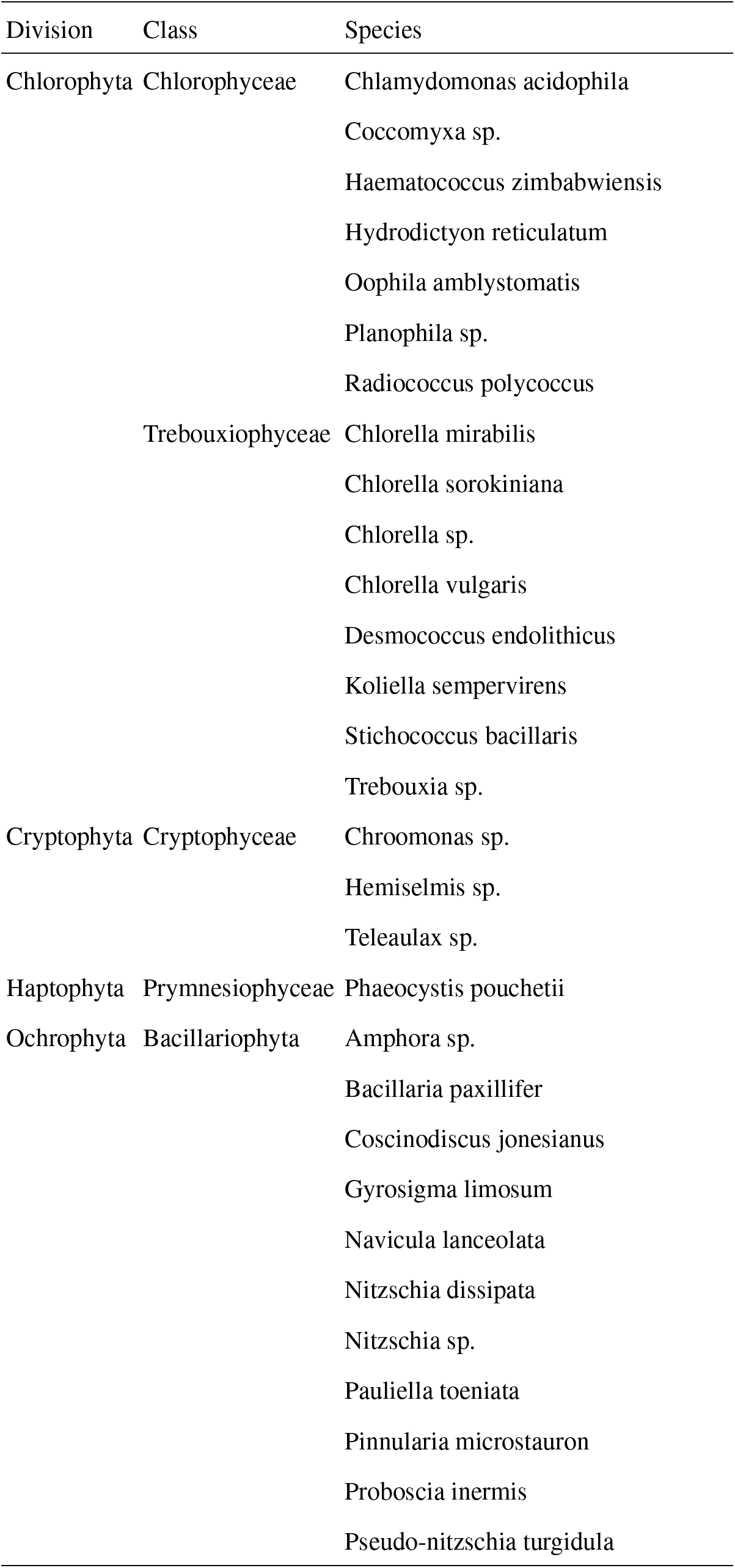
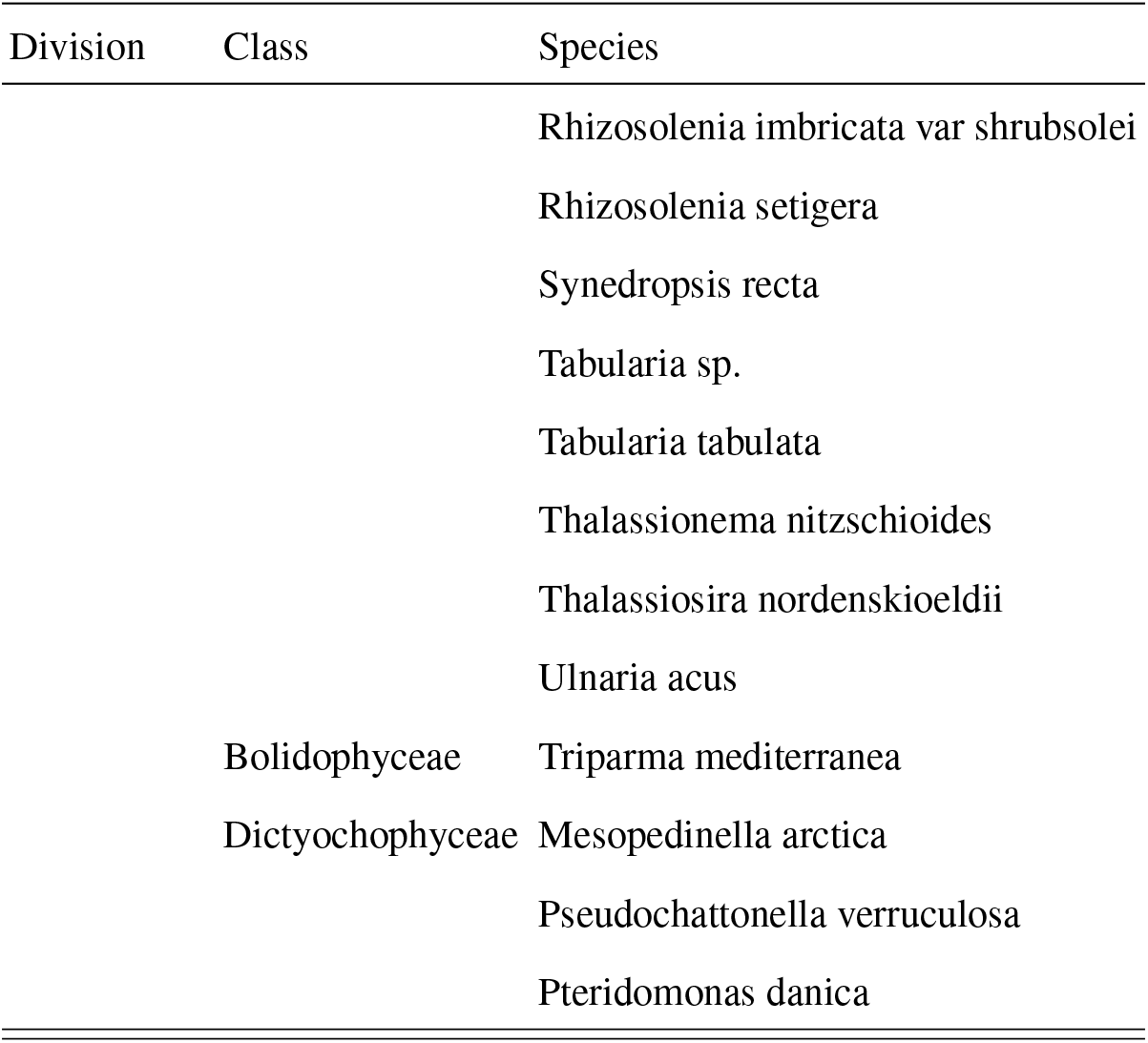
List of species in the metabarcoding data sets only found in the deep samples (from 10 to 50 m).

**Table S3.**
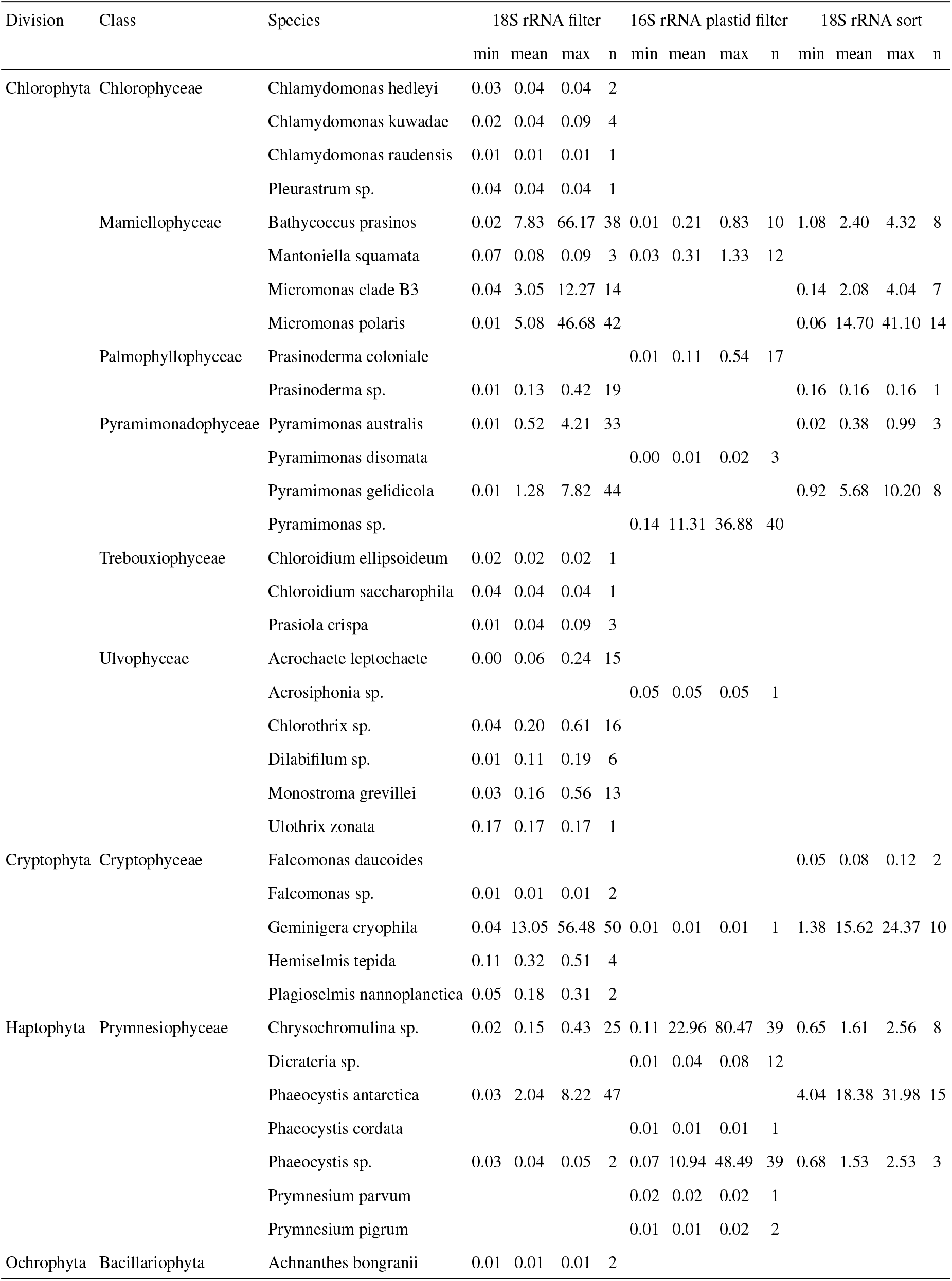
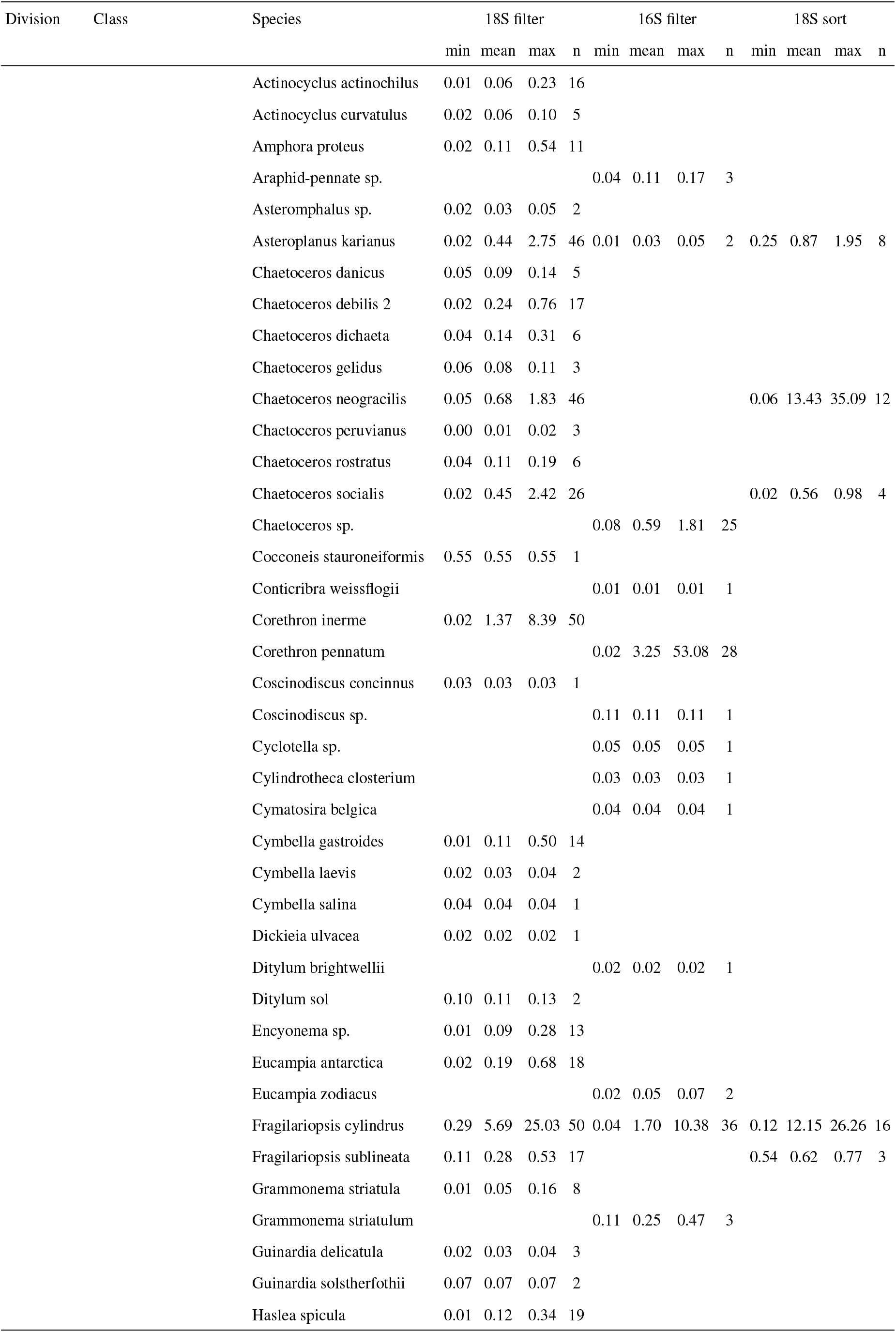
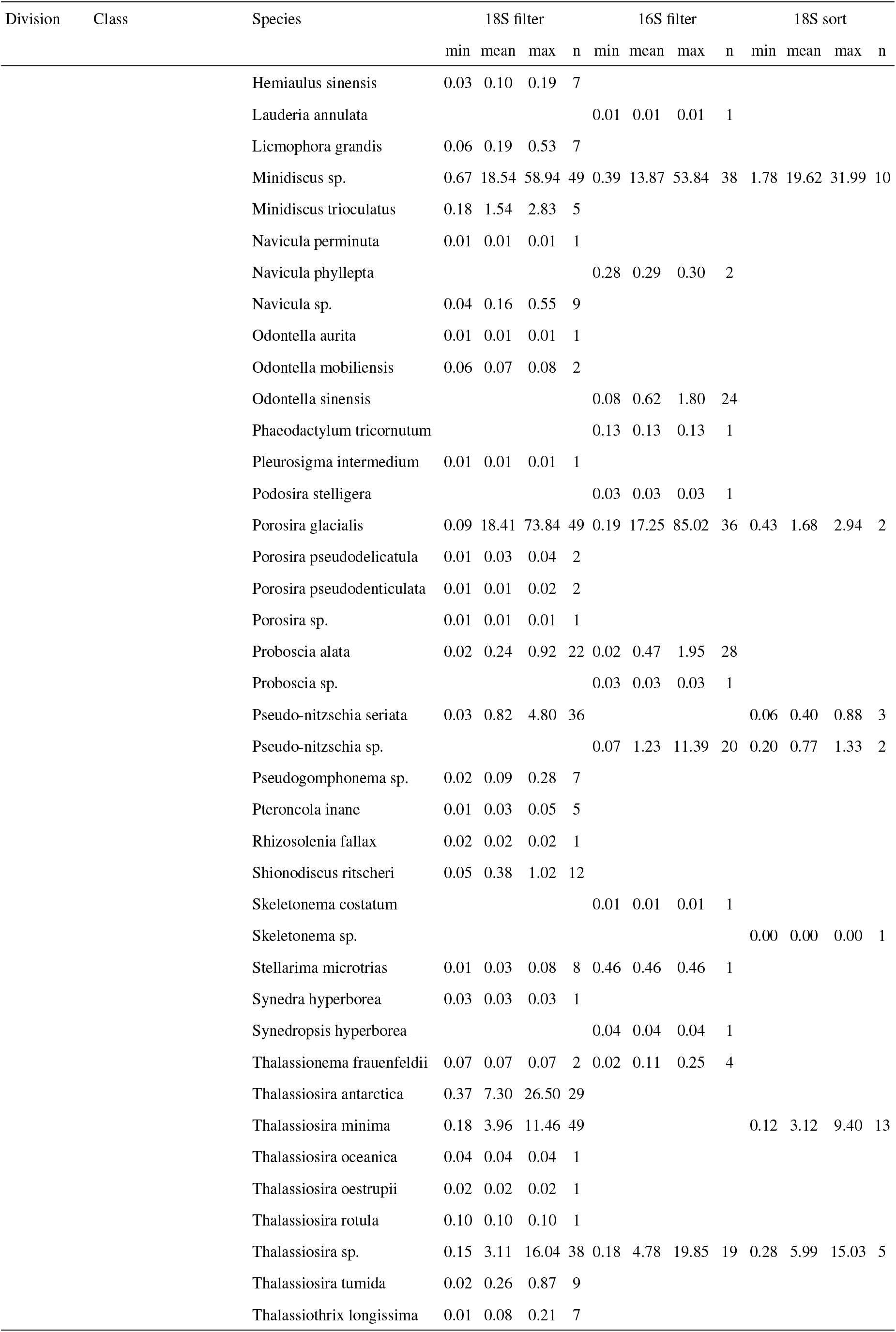
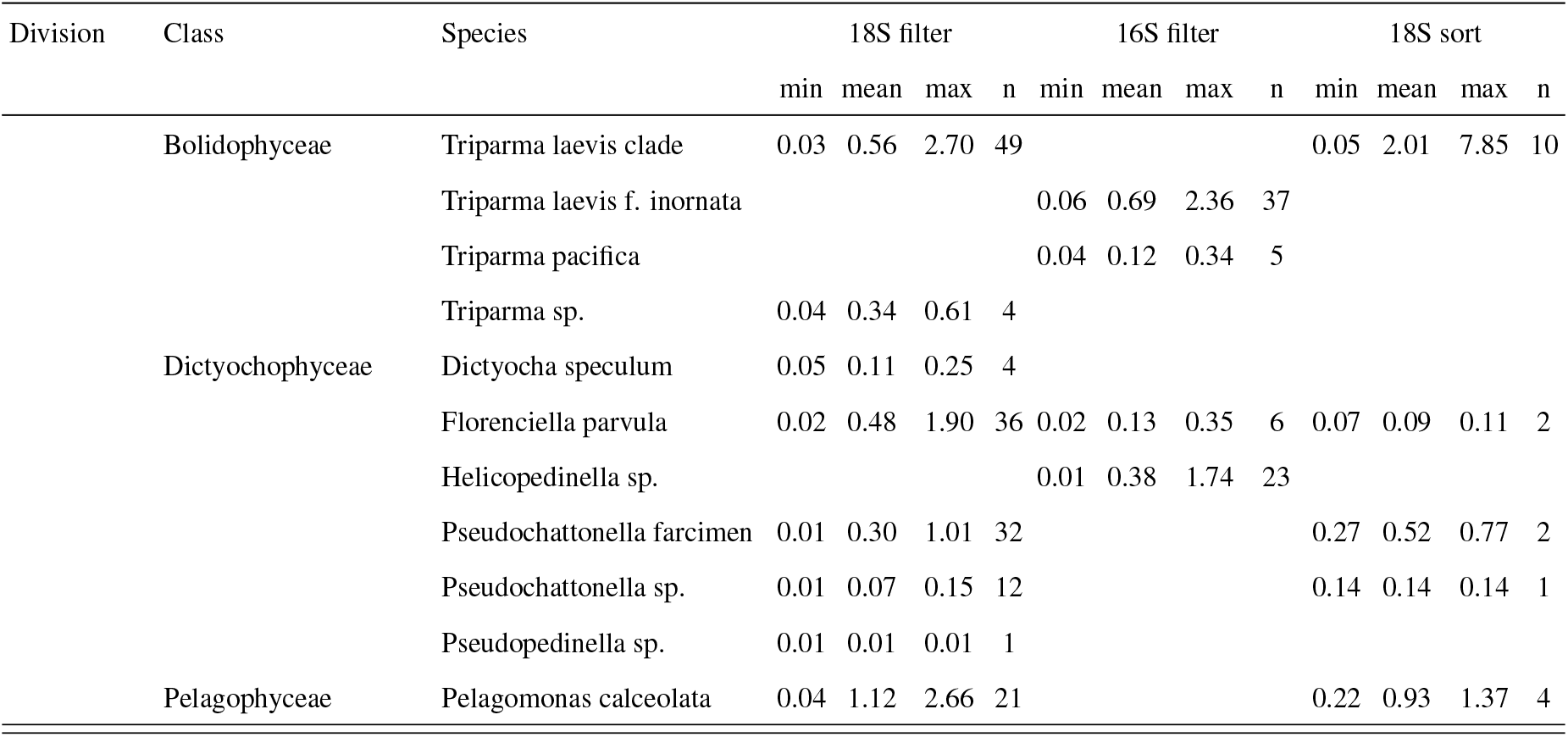
List of species found in the metabarcoding data sets for the surface samples. Minimum (min), mean (mean) and Maximum (max) contribution (in %) to the photosynthetic metabarcodes and the number of samples (n) where found for the 18S-filter, 16S-filter and 18S-sort datasets.

**Table S4.**
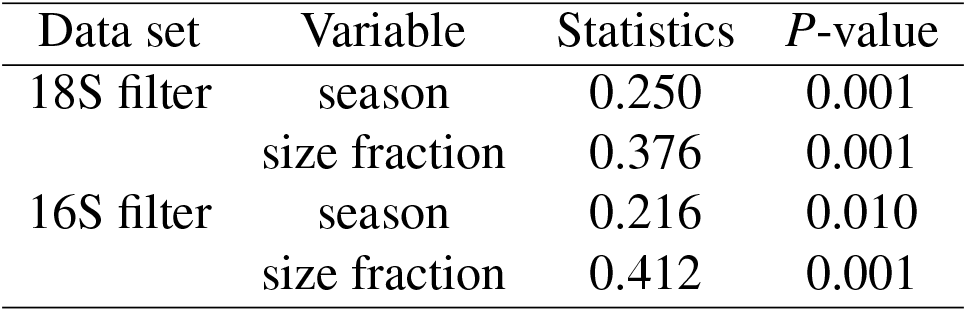
ANOSIM analysis for surface samples contrasting the effect of season or size-fraction.

**Figure S1.**
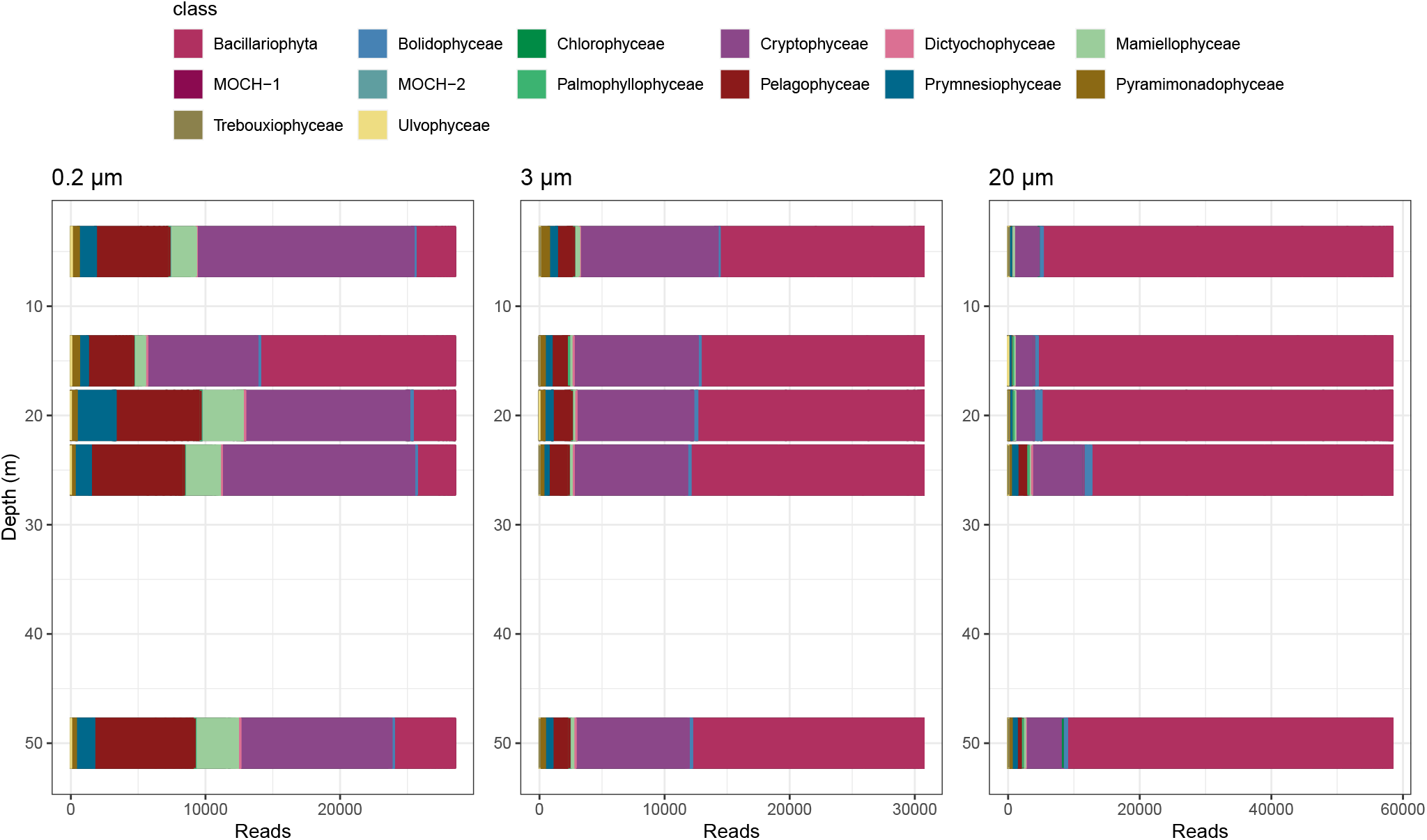
Community composition of phytoplankton at the class level along a vertical profile obtained on January 16, 2015, from 5 m and down to 50 m, based on the 18S rRNA gene for filtered samples.

**Figure S2.**
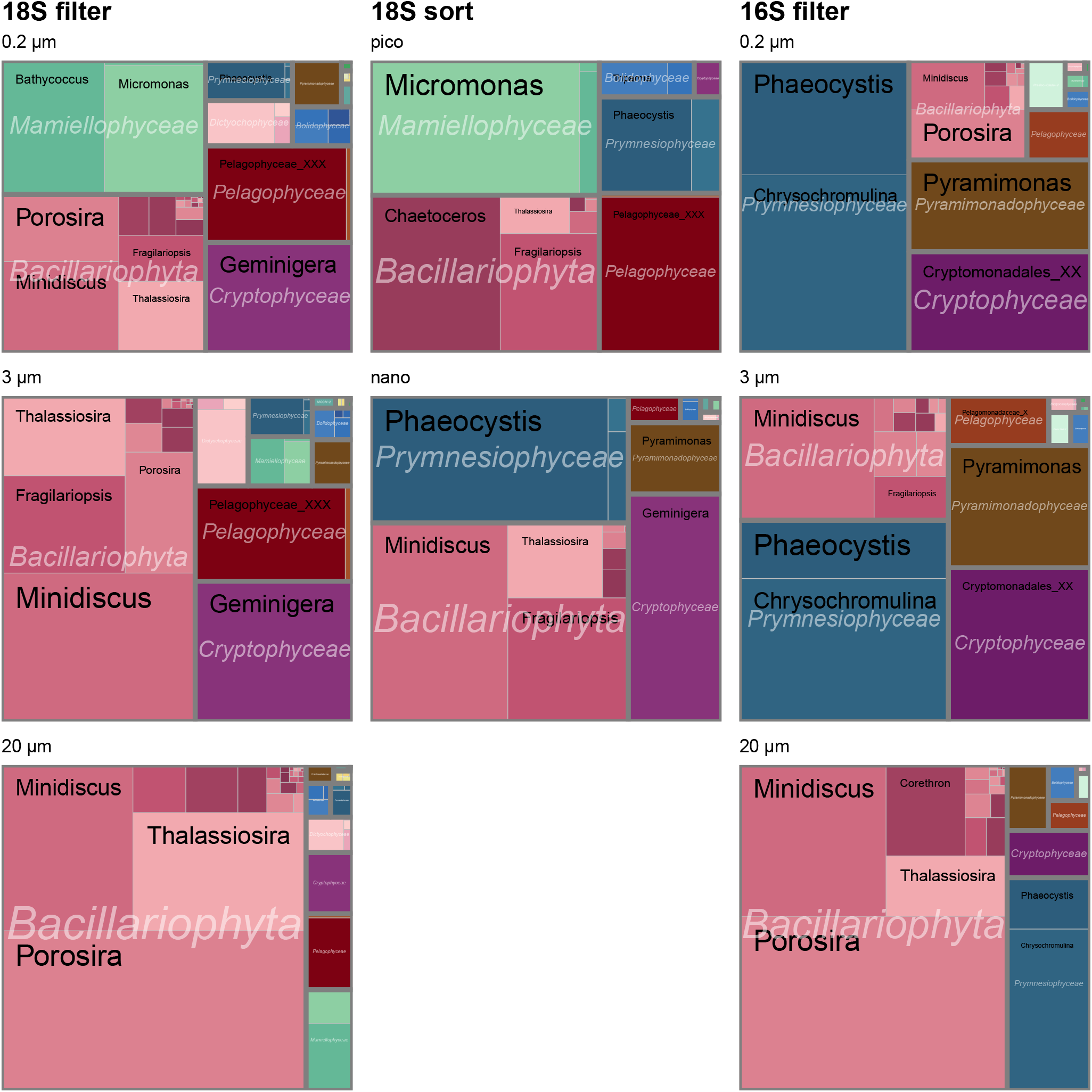
Relative abundance of the different genera in surface samples based on three different amplicon sequencing approaches for each size fraction. Left: 18S rRNA gene on filtered samples. Middle: 18S rRNA gene on sorted samples. Right: plastidial 16S rRNA gene on filtered samples.

**Figure S3.**
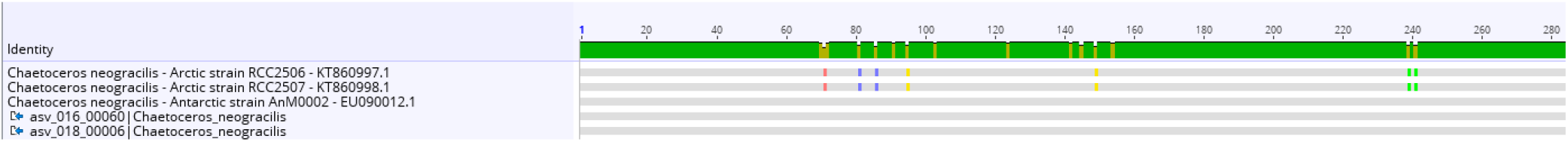
Sequence alignment of 18S rRNA ASVs for *Chaetoceros neogracilis* showing the differences between Arctic and Antarctic strains sequences. The ASVs from this study are identical to the Antarctic strain and show 7 bp differences to Arctic strains.

**Figure S4.**
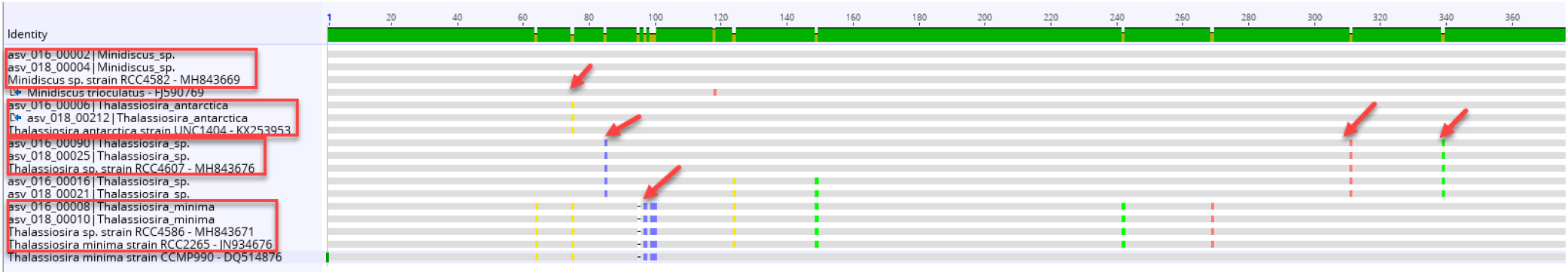
Sequence alignment of 18S rRNA ASVs for major *Thalassiosira* and *Minidiscus* ASVs in comparison to reference sequences.

**Figure S5.**
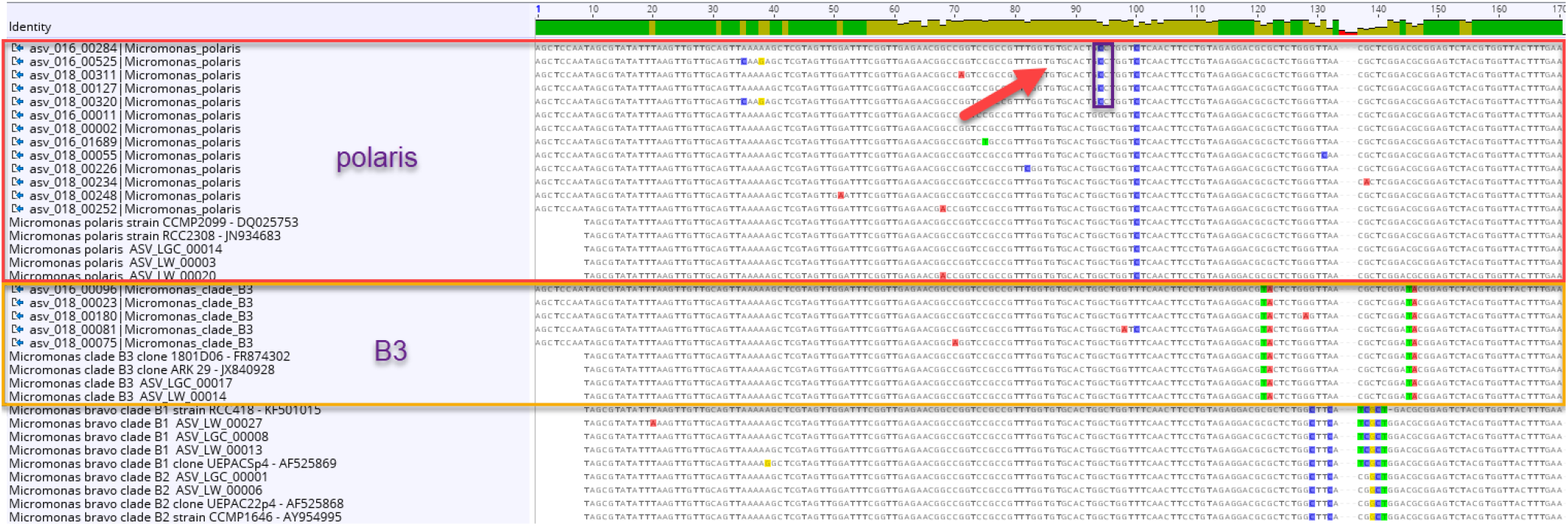
Sequence alignment of 18S rRNA ASVs for *Micromonas* showing the clear signatures for *M. polaris* and clade B3 (Tragin and Vaulot 2019). Within *M. polaris* some sequences have a different signature pointing to a new clade specific of Antarctic waters (arrow).

**Figure S6.**
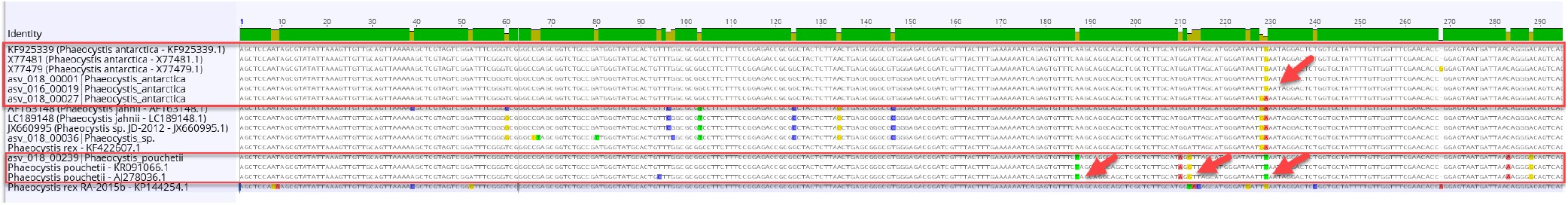
Sequence alignment of 18S rRNA ASVs for *Phaeocystis* showing the clear signatures for *P. antarctica* and *P. pouchetii*.

**Figure S7.**
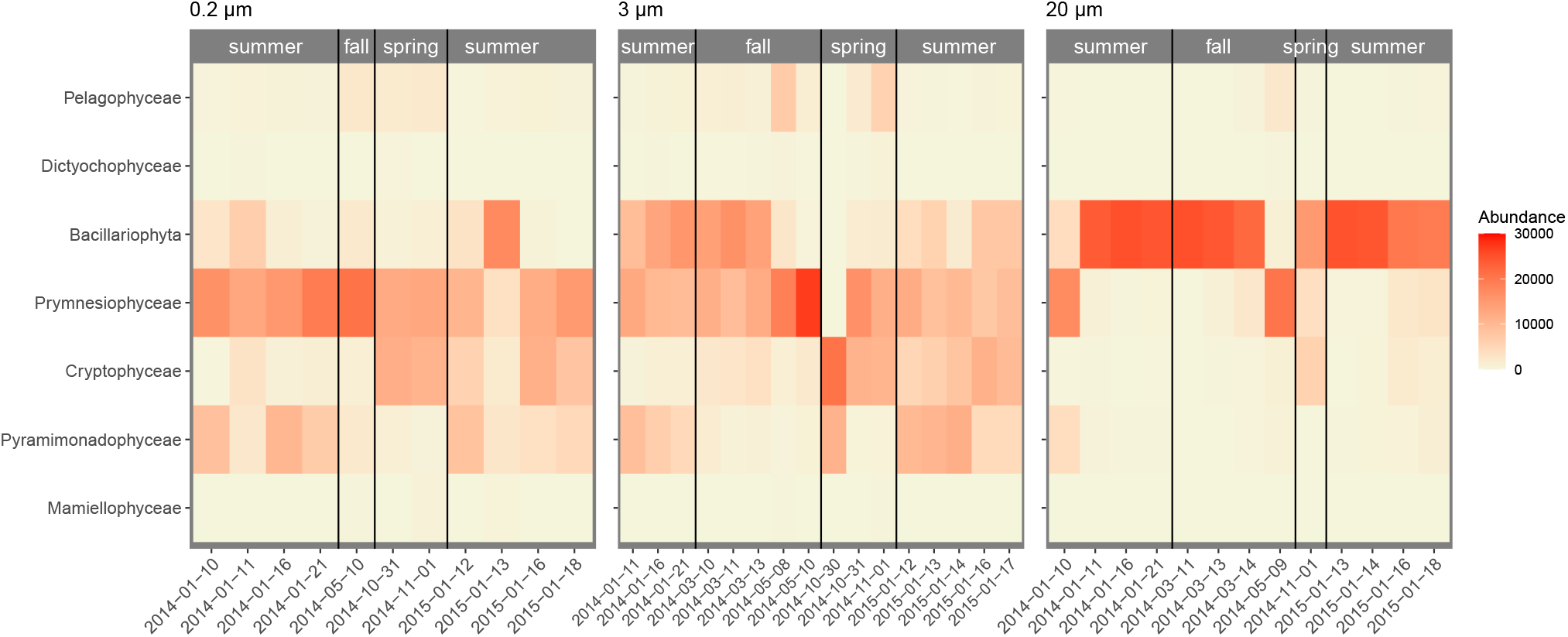
Read numbers for the main photosynthetic taxonomic groups at the class level for plastidial 16S rRNA gene for filtered surface samples. The color scale of the heatmap corresponds to the normalized number of reads of each taxon. Season delimitation corresponds to meteorological seasons. Left: 0.2-3 μm. Middle: 3-20 μm. Right: > 20 μm.

**Figure S8.**
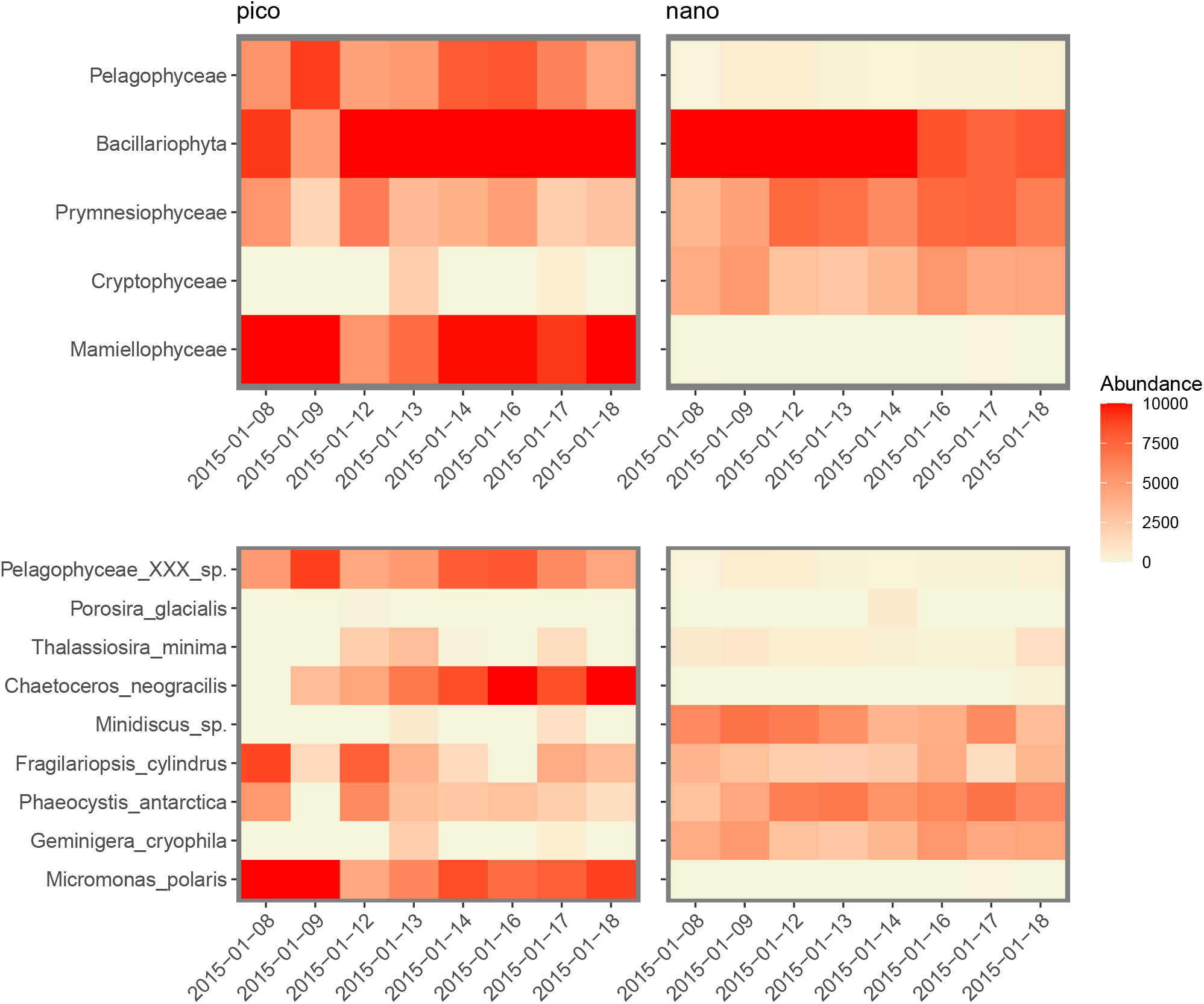
Read numbers for the main photosynthetic taxonomic groups at the class (Top) and genus (Bottom) levels of 18S rRNA gene for sorted samples from surface waters. The color scale of the heatmap corresponds to the normalized number of reads of each taxon. Left: pico size fraction. Right: nano size fraction.

**Figure S9.**
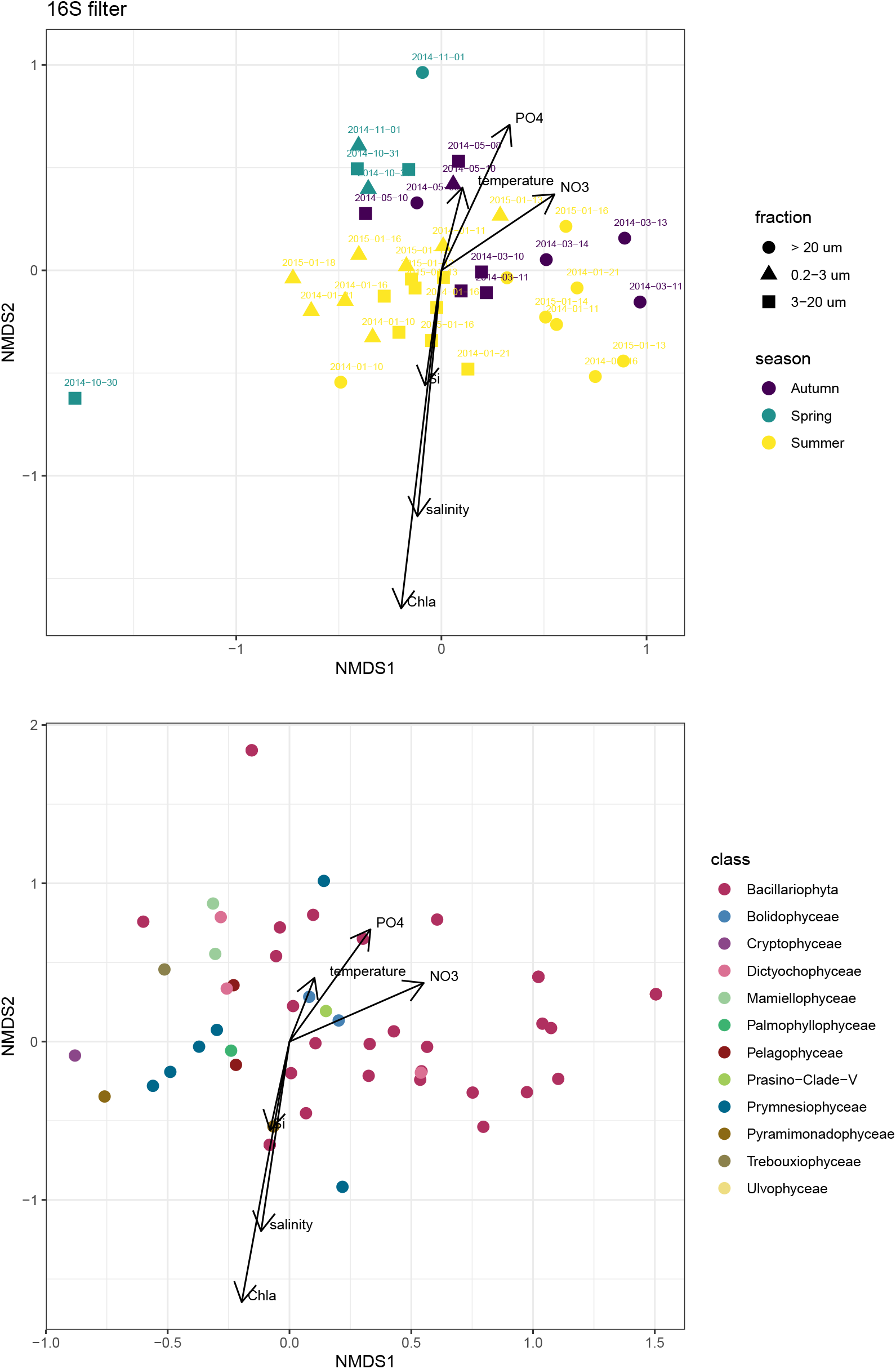
Non-metric multidimensional scaling (NMDS) analysis based on Bray-Curtis dissimilarities of the phytoplankton community composition (species) labeled by meteorological season and size fraction using the plastidial 16S rRNA gene. (A) Samples. (B) ASVs. Stress = 0.15.

